# Cortex-wide voltage imaging in a sensory cued reaching task reveals fast subnetwork dynamics

**DOI:** 10.1101/2025.04.24.650533

**Authors:** Yunmiao Wang, Soon Ho Kim, Ellie Jiayi He, Brune Le Chatelier, Hannah Choi, Dieter Jaeger

## Abstract

Oscillatory activities and network dynamics are fundamental to sensorimotor processing, motor control, and cognitive functions. Capturing the fast cortical dynamics underlying sensorimotor behaviors demands temporal resolution beyond conventional imaging methods. We employed pan-cortical JEDI-1P voltage imaging of layer 2/3 activity in a skilled reaching task to reveal previously inaccessible rapid networks dynamics. Our approach uncovered distinct spectral signatures including reward-related gamma oscillations in M2, global low-frequency suppression during movement execution, and 8 Hz large-amplitude oscillations in sensorimotor cortex during task disengagement. Using independent component analysis, we were able to separate multiplexed functional networks with fast temporal features shared among sensorimotor regions and slower dynamics such as a global preparatory ramp. These distributed activity patterns successfully predicted both sensorimotor parameters and trial outcomes. Our study bridges critical gaps in understanding rapid temporal organization of cortical networks during complex behaviors, revealing how fast neural dynamics coordinate to produce motor actions across distributed cortical regions.

## Introduction

Brain network oscillations have long been proposed as a mechanism of coordinating dynamic routing of information processing in the brain^1, 2^. Oscillatory activity of different frequency bands is associated with various sensory, motor, and cognitive functions in primates^3, 4^ as well as rodents^5, 6, 7^. Beta oscillations (13–30 Hz) in the cortico-basal ganglia pathway play a role in sensorimotor processing^8^ and are also linked with suppression of movement initiation^9^. While beta oscillations mainly affect motor functions, synchronized gamma oscillations of 30–80 Hz are proposed to influence attention^10, 11, 12^, sensory processes^13, 14, 15^, and working memory^16^. Slow-wave activity such as theta (4–8 Hz) and alpha or mu oscillations (8–13 Hz) can also influence the multisensory processing and network-wise communication in the brain^17, 18, 19, 20^ and are generally suppressed during active movements^21^.

Fast cortical processing beyond the temporal resolution of calcium imaging is not limited to oscillations, as neural dynamics related to sensory events and motor control also evolve with millisecond precision. Substantial work employing widefield calcium imaging in head-fixed mice has revealed rich pan-cortical network dynamics in relation to behavior^22, 23, 24, 25, 26, 27^. This work has been predominantly focused on layer 2/3 in cortex due to ease of access for imaging but also rich network dynamics in this layer due to widespread lateral connections. L2/3 pyramidal cells are intratelencephalic neurons projecting in a local network to other L2/3 neurons and L5 pyramidal cells, as well as widely to other cortical areas, connecting sensory to motor processing, and finally to striatum but not other subcortical targets^28, 29^. Importantly these connections include callosal fibers connecting to the opposite hemisphere^30^ allowing for interhemispheric communication. Functionally, L2/3 neurons are part of recurrent network activity that undergoes synaptic plasticity during learning^31, 32^. Even a small number of L2/3 neurons in motor cortex, when stimulated, can trigger training-specific motor dynamics^33^. L2/3 usually shows sparse pyramidal neuron activity due to strong local PV type interneuron activity^34^ and that interaction can also form the basis of a pronounced gap-junction dependent 30-45 Hz gamma rhythm^35^. An important gap in our understanding of L2/3 network dynamics concerns the question of fast temporal dynamics of identified networks in relation to behavior.

In this study, we introduce widefield pan-cortical L2/3 voltage imaging in head-fixed mice as a novel method to discern the fast dynamics of cortical networks underlying sensorimotor integration during a cued water-reaching task. The reinforced reaching task effectively assesses cortical network processing involved in cue evaluation, motor preparation, skilled movement execution, and reward uptake^36, 37, 38^. Our study reveals that all aspects of the task engage widespread cortical networks that can be further separated into subnetworks with distinct temporal dynamics that link them to task processing. Gamma-band mu oscillations were found to be engaged in task-specific roles. Independent component analysis revealed fast and slow temporal features in sensorimotor regions and global cortical networks. Further, the expression of these networks in single trials allows the prediction of behavioral performance.

## Results

### High-frequency oscillations accompany a sensory cued reaching task

To explore fast cortical network dynamics during behavior, we performed high-speed wide-field voltage imaging during a sensory-guided reaching task. We opted to use JEDI-1P, a recently developed voltage sensor^39^, which enables recording through an intact skull at 200 Hz and tracking of population neural activity up to 100 Hz (**Fig.1a and Methods).** Animals were trained to perform a directional water reaching task **(Fig. 1b)** that was previously demonstrated to depend on cortical activity^38^. In each trial, a gentle train of air-puff stimuli on either left or right whiskers was followed by a delay period with variable length of 0.75 to 1.25s. After a visual go-cue, a droplet of flavored sucrose water was delivered. For four out of the seven animals, the reward was delivered on the same side as the whisker stimulus; for the remaining animals, it was delivered on the opposite side. The animals were trained to reach for the reward with the limb on the same side as the water droplet during the response period (**Fig. 1b-c**). Snapshots of whole-cortex artifact-corrected –ΔF/F revealed widespread cortical activation before and during reaching (**Fig. 1d**), as was previously found with calcium imaging methods^37, 40^. Unlike calcium imaging that reports calcium concentration as a proxy of neural activity, voltage sensors with a wide dynamic range such as JEDI-1P possess a unique advantage of tracking subthreshold activity via a direct measurement of voltage changes on the neuronal membrane. Moreover, the fast voltage response of this sensor allows resolving of oscillations up to the gamma range^39^. Our initial observations of frequency specific intrinsic signaling revealed an increase in gamma power in M2, which started at reward delivery and lasted until after the reach movement **(Fig. 1e)**. This distinct mouse frontal M2 gamma network is described here for the first time to our knowledge. Gamma activity in relation to motor control has previously been observed in single unit activity in SMA of primates^41^ and human ECoG and LFP M1 recordings^42^ where it is related to learning. In addition to peri-movement gamma oscillations, we observed a cortex-wide suppression of low-frequency power that lasted through the entire trial starting with the air-puff cue (**Fig. 1e**). This low-frequency suppression is reminiscent of event related EEG desynchronization^21^ reflecting an active brain state^43^.

**Figure 1.**
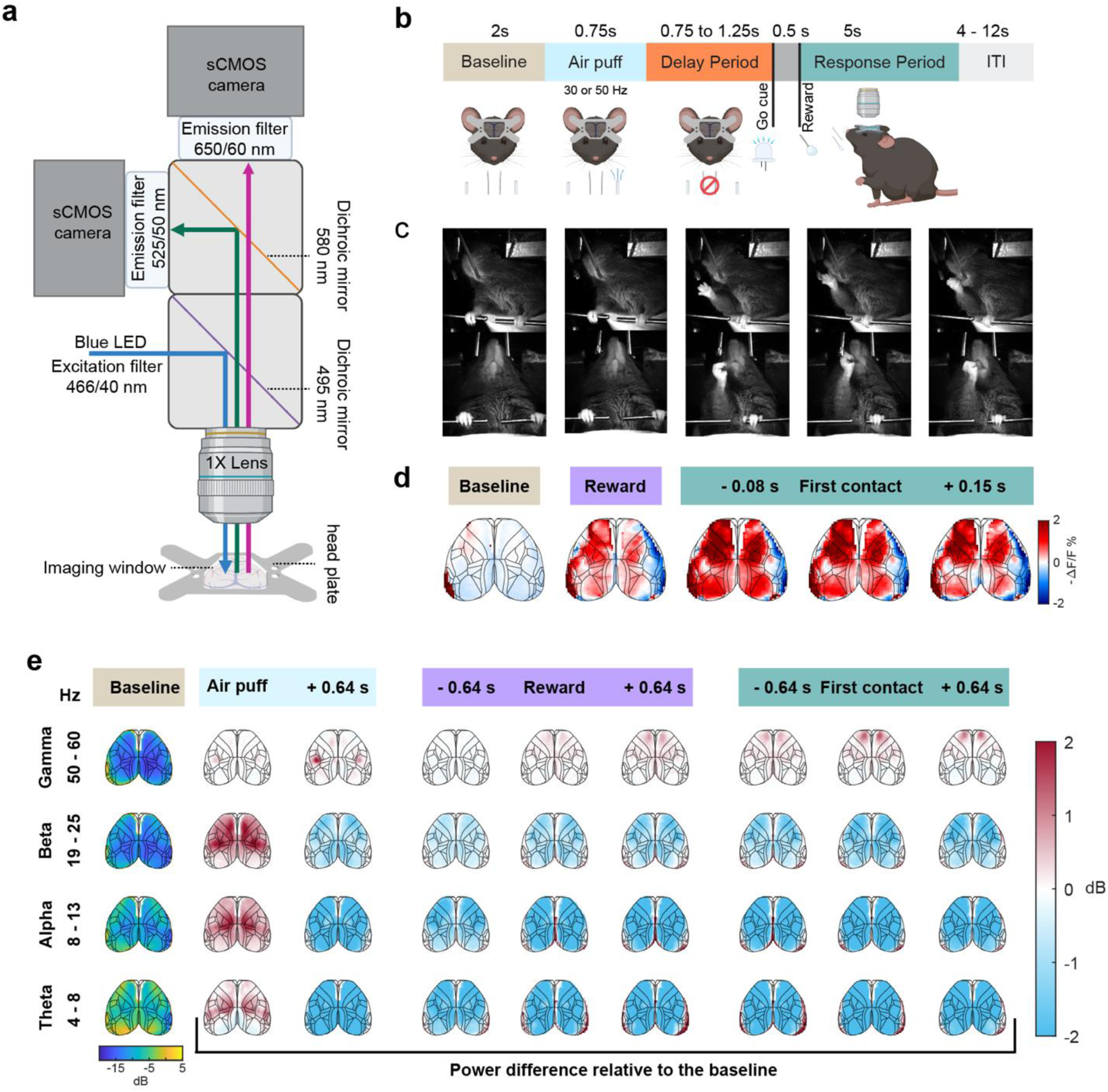
High- and low-frequency spectral changes accompany a sensory cued reaching task. (a) Imaging setup schematic. The voltage sensor in green, JEDI-1P, and the reference fluorescent in magenta, mCherry, were excited by the same blue LED light source. The emission light was captured by two CMOS cameras with a field of view of 10 × 10 mm^2^. All imaging data in this study was acquired at 200 Hz. (b) Design of the bilateral reaching task. Animals were presented with two air-puff lines that applied gentle stimuli to either side of the whiskers and with two waterspouts that delivered sucrose reward. A trial started with a baseline period and followed by air-puff train stimuli at either 30 Hz or 50 Hz. After a delay period with variable length, a visual cue was on for 0.5 s before reward was delivered on the side associated with the whisker stimuli. Animals were trained to reach for the reward with the limb on the same side as the reward. (c) Snapshots of video recording during a successful trial demonstrating the reach for reward. Panels from left to right indicate baseline, reward presence, reaching, first contact with reward, and reward retrieval, respectively. (d) Spatial maps of single-frame average –ΔF/F aligned with the time points shown in (c) during right-side reaches of success trials. A representative animal of 7 mice, n = 941 trials for baseline and reward periods and n = 924 for response period. (e) Bilateral increase in gamma activity occurred during the reward period of success trials. Low frequency activity was suppressed following the sensory stimuli and intensified during the reaching. Left most column shows baseline power of each frequency range. The spatial maps in blue, white and red represent differences of power between an event aligned window and the baseline at a given frequency range. Each spatial map is aligned to the Allen Atlas. n = 7 mice. Note an additional stimulus driven gamma peak during the air-puff train that was delivered at 50 Hz to right whiskers and 30 Hz to left whiskers.

### Oscillatory network engagement is dependent on task performance

Next, we addressed the question if these intrinsic spectral changes were related to specific behavioral states by comparing power changes in different frequency ranges between left- and right-reaching trials and between success trials and no-response trials.

Trials in which reward was retrieved and consumed during the response period were labeled as success trials, and those in which reward was delivered but not retrieved during the response period were labeled as no-response trials. We defined the response time as the time between reward delivery and the first contact of the reaching forelimb with the reward spout (**Fig. 2a**). The median response time of success trials was 1.17 s and 1.13 s for left-side and right-side trials respectively (**Fig. 2b)**. Peri-event activity of individual trials was used to calculate frequency specific power changes during pre-trial baseline and different task events, including air-puff stimulation period, reward period, and response period (**Fig. 2c**). We first compared baseline-subtracted power in each frequency band between left-side and right-side success trials to evaluate potential lateralization of neural oscillations. The comparison revealed higher gamma power in M2 on contralateral side to the reaches during the response period (**Fig. 2d, e)**. Intriguingly, most other spectral changes did not appear to be different for left- and right-reaching trials, revealing an overall bilaterally symmetric task-processing mode (**Fig. 1e and Supplementary Fig. 1**). Differences in the responses between success and no-response trials were then assessed in each frequency band (**Fig. 2f-h**). The spatial maps of power differences highlighted that a bilateral gamma increase in M2 was uniquely present in success trials. In contrast, the task-related suppression at low frequencies (alpha, theta, low beta) was more pronounced in success trials globally, especially at sensorimotor regions (**Fig. 2g and Supplementary Fig. 1**). The spectral power changes over time at regions of interest (ROIs) further showed an increasing divergence of signal power during no-response trials starting well before the reaching movement (**Fig. 2h)**.

**Figure 2.**
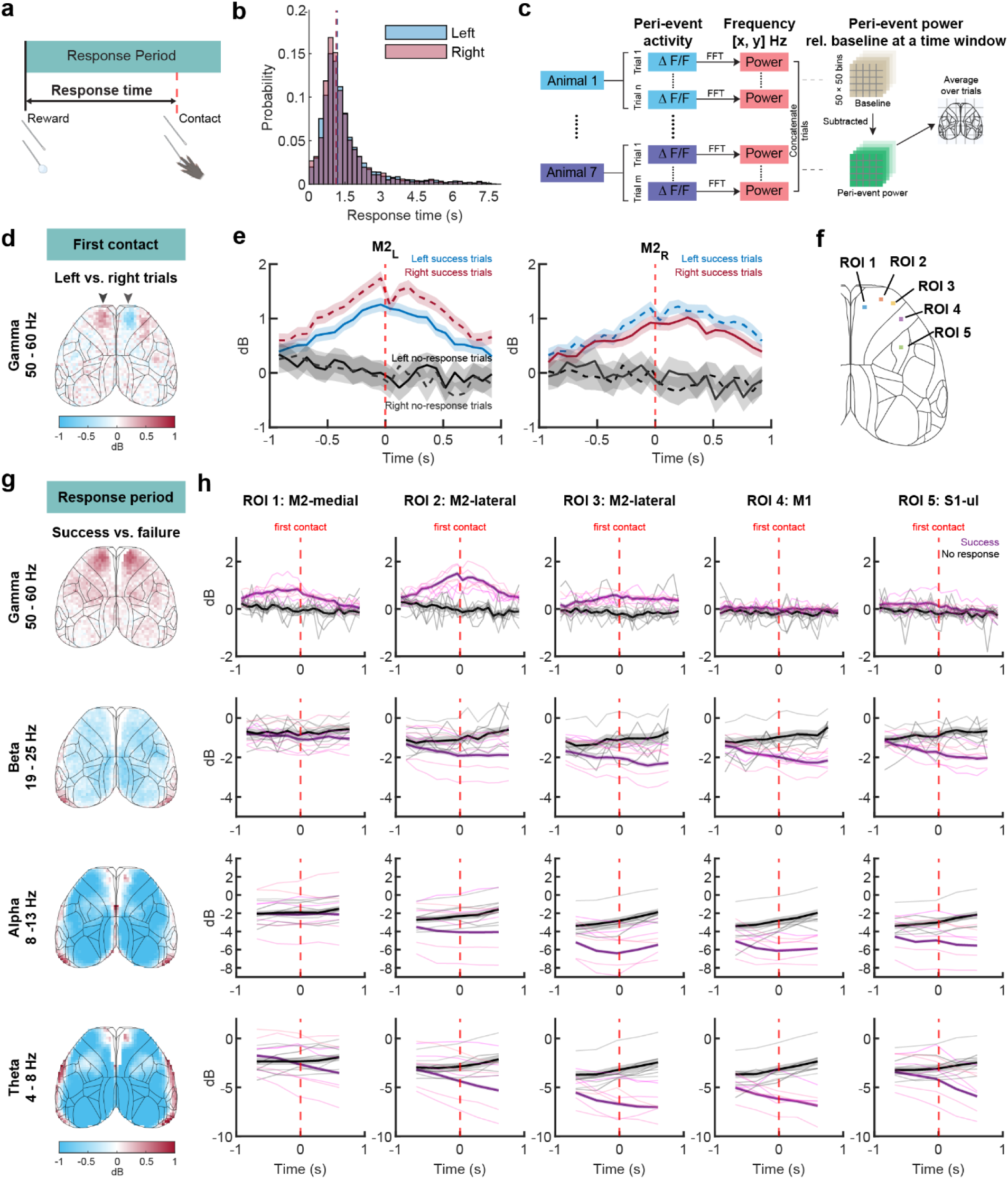
Spectral power diverges based on trial outcomes. (a) Schematic shows the definition of response time. (b) The distribution of response time in success trials guides selection of window size. Vertical lines indicate the median time for left and right trials, 1.17 s and 1.13 s, respectively. n = 3346 trials for left trials, and n = 3262 for right trials. P value = 0.0258, Two-sided Mann-Whitney U Test on the response time between left- and right-trials. (c) Schematic shows how changes of power at a given frequency range were calculated. Power was calculated based on peri-event activity aligned with specific events including air-puff onset, reward onset, and first contact with waterspout. (d–e) M2 contralateral to the reaching limb shows stronger gamma power at 50 – 60 Hz range than ipsilateral M2. (d) The spatial map represents difference in power between left- and right-reaching success trials. n = 7 mice. (e) Time courses of power at bilateral M2 during left- and right-reaching success trials and no-response trials. The power of success trials is calculated based on 2 s activity centered around first contact, indicated with vertical dotted red line, and that of no-response trials is centered around 2 s following the reward delivery. n = 3241, 876, 3179, and 753 trials, for left-reaching success, left-reaching no-response, right-reaching success, and right-reaching no-response, respectively. (f) Regions of interest (ROIs) used in h. Each ROI is 2 × 2 pixels (0.2 × 0.2 mm^2^). (g) Power differences between success and no-response trials demonstrated that the increase in gamma power and suppression in beta power were stronger in success trials. Spatial map of the power difference is aligned with the moment of first contact during the response period. N = 7 mice. (h) Spectral changes diverged significantly between success and no-response trials during the response period, with higher gamma increase at M2 and more low-frequency suppression globally in success trials. Red vertical dashed line indicates the moment of contact with reward. n = 6420 success trials and 1629 no-response trials. Lines in (e) and (h) indicate mean baseline-subtracted power across trials from all mice, and shaded area denotes 95% CI.

### Movement is suppressed during bilateral ∼8Hz oscillations

Spectral power (**Fig. 3a-d**) and distinct oscillatory periods around 8 Hz were notably more prevalent in no-response trials than success trials. While single trial –ΔF/F traces from bilateral ROIs showed large-amplitude 8Hz oscillations in both success and no-response trials, these oscillations were detected in 12.4 % of no-response trials but only in 1.5 % of success trials (**Fig. 3c, e**). When trials were grouped based on the presence or absence of these large-amplitude 8Hz oscillations during the response period, grouped trials with such oscillations present showed increased power bilaterally at the frontal cortex and V1 irrespective of the trial outcomes (**Fig. 3d, f, g** middle row). 8Hz oscillations of no-response trials occurred sporadically during the entire trial, whereas in success trials they subsided by the time the mice started reaching (**Fig. 3f-g**). Concurrent limb speed was significantly lower when 8Hz oscillations were present (**Fig. 3f-h**). In fact, when aligning paw position traces with these oscillations in individual trials, a complete cessation of paw movement during 8Hz oscillations was observed (**Fig. 3e**). This finding reveals a new antikinetic role of 8Hz-oscillations in mouse cortex and indicates that they are likely related to task-disengaged brain states previously described^44^. To further relate 8Hz-oscillations to the arousal level of the mouse, we determined if pupil diameter changed during 8Hz-oscillatory activity. We found that no-response trials with detected 8Hz oscillations showed larger pupil diameter early in each trial compared to the rest of the groups (**Fig. 3i**), suggesting an elevated arousal level associated with the oscillations despite the absence of movement.

**Figure 3.**
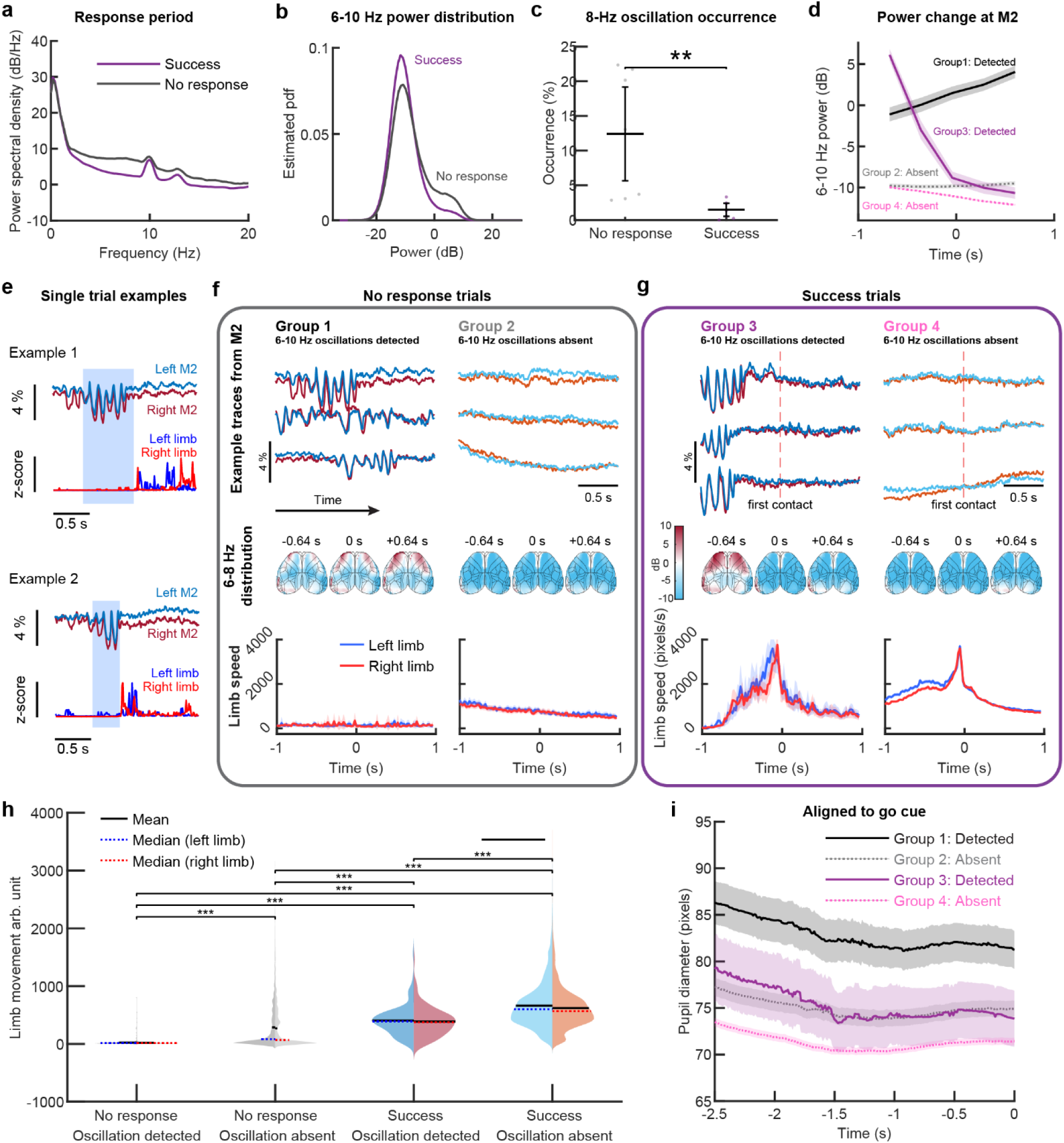
Movement is suppressed during bilateral 8Hz oscillations. (a–b) 6–10 Hz power is higher in no-response trials than that in success trials during the response period. –ΔF/F from bilateral M2, the center of 8-Hz oscillation occurrence, is used for the spectral analysis. N = 1620 no-response trials and 6370 success trials. (a) Power spectral density between 6-10 Hz from success trials is higher than that from no-response trials. (b) Distribution of power between 6-10 Hz is higher in no-response trials than that of success trials. (c) 8-Hz oscillations with large amplitude are observed more frequently in no-response trials (12.4 %) than success trials (1.5 %). Two-sided Mann-Whitney U Test was performed on the oscillation occurrence between two trial outcomes (N = 7 mice). P value is 0.0023. Error bar denotes 95% CI. (d) Power between 6–10 Hz shows significant differences when success and no-response trials are grouped based on the detection of oscillations. Gray color indicates no-response trials (groups 1 and 2), and magenta and purple indicate success trials (groups 3 and 4). Solid and dashed lines indicate the mean power change from M2 in trials exhibiting 8 Hz oscillations (groups 1 and 3) and in those that do not (groups 2 and 4), respectively. Shaded areas denote 95% CI. N = 263, 1357, 113, and 6257 trials, respectively. (e) Movements are absent when large-amplitude oscillations occur demonstrated with example –ΔF/F traces from bilateral M2 and concurrent movement (f-g) The 8-Hz oscillations mainly happen at the frontal part of the brain. Limb movements are typically absent during these oscillations. (f) Typical neural activity and limb speed with and without 8 Hz oscillations in no-response trials. Top subpanel within the gray box shows example -ΔF/F traces from M2, highlighting the presence or absence of 8-Hz oscillations. Middle panel shows the 8-13 Hz power across the cortex over time within each group. Bottom panel shows concurrent limb speed. N = 263 trials where 8-Hz oscillations were detected, and 1357 trials where they were not detected. (g) Same as (f) for success trials. (h) The amount of limb movement is significantly lower when 8 Hz oscillations occurred. Each violin plot is split into two halves, where the left side represents left limb movement, and the right side represents the right limb movement. The median limb speed from individual trials was used to represent the amount of movement in each trial. Solid horizontal line denotes the mean speed of each given distribution, and dashed line denotes median trials. Two-sided Mann-Whitney U-test was run between each group of speed of the same limb. P values are smaller than 10^-14^ for all twelve paired tests. (i) Pupil diameter from no-response trials with 8-Hz oscillations is significantly larger than that in other groups even prior to the Go cue. N = 235, 1160, 101, and 5031 trials in Group 1-4, respectively, from 4 mice. Pupil data was not acquired in the other 3 mice.

### Low-dimensional spatiotemporal activity dynamics related to sensorimotor processes are revealed by independent component analysis

Given the success of dimensionality reduction methods in extracting low-dimensional cortical activity manifolds related to motor control from single unit data^45, 46^ and calcium imaging^47, 48, 49^, we set out to examine how dimensionality reduction can be applied to widefield voltage imaging to assess cortical network dynamics. Temporal independent component analysis (t-ICA) decomposes high-dimensional time series into temporal components that are assumed to be statistically independent^50^. This method has been applied broadly on neural data, including fMRI^51^ and EEG signals^52^, and can be used to identify temporally distinct components of the dynamics^53, 54, 55, 56, 57^. To separate networks with distinct dynamics we employed temporal ICA (t-ICA) to decompose the temporal signals into non-Gaussian and optimally independent network dynamics (**Fig. 4a**). To validate the t-ICA method, we first applied it on peri-event activity aligned with the onset of 30-Hz air-puff stimuli on the left whiskers (**Fig. 4b-c**) where we know the ground truth of an initial contralateral barrel cortex activation. Generally, six independent components (ICs) were sufficient to reduce the 1400-dimensional pixel space to distinct temporal dynamic features that can reconstruct the signal with correlation coefficient above 0.93 on testing datasets (**Fig. 4d-g**). The decision of using six ICs was made based on the commonly used elbow method in machine learning by varying the number of ICs (**Fig. 4e, f**). We chose the minimal number of ICs that adequately represent the original signal by quantifying the similarities between the original and reconstructed signals (**Fig. 4e**) while best satisfying the condition of non-Gaussian distribution of the ICs by measuring their mean absolute kurtosis (**Fig. 4f**). The same validation process shows that 6 components were sufficient to reconstruct for other peri-event activity aligned with the reward delivery and first contact with the waterspout delivering reward (**Fig. 4g**).

**Figure 4.**
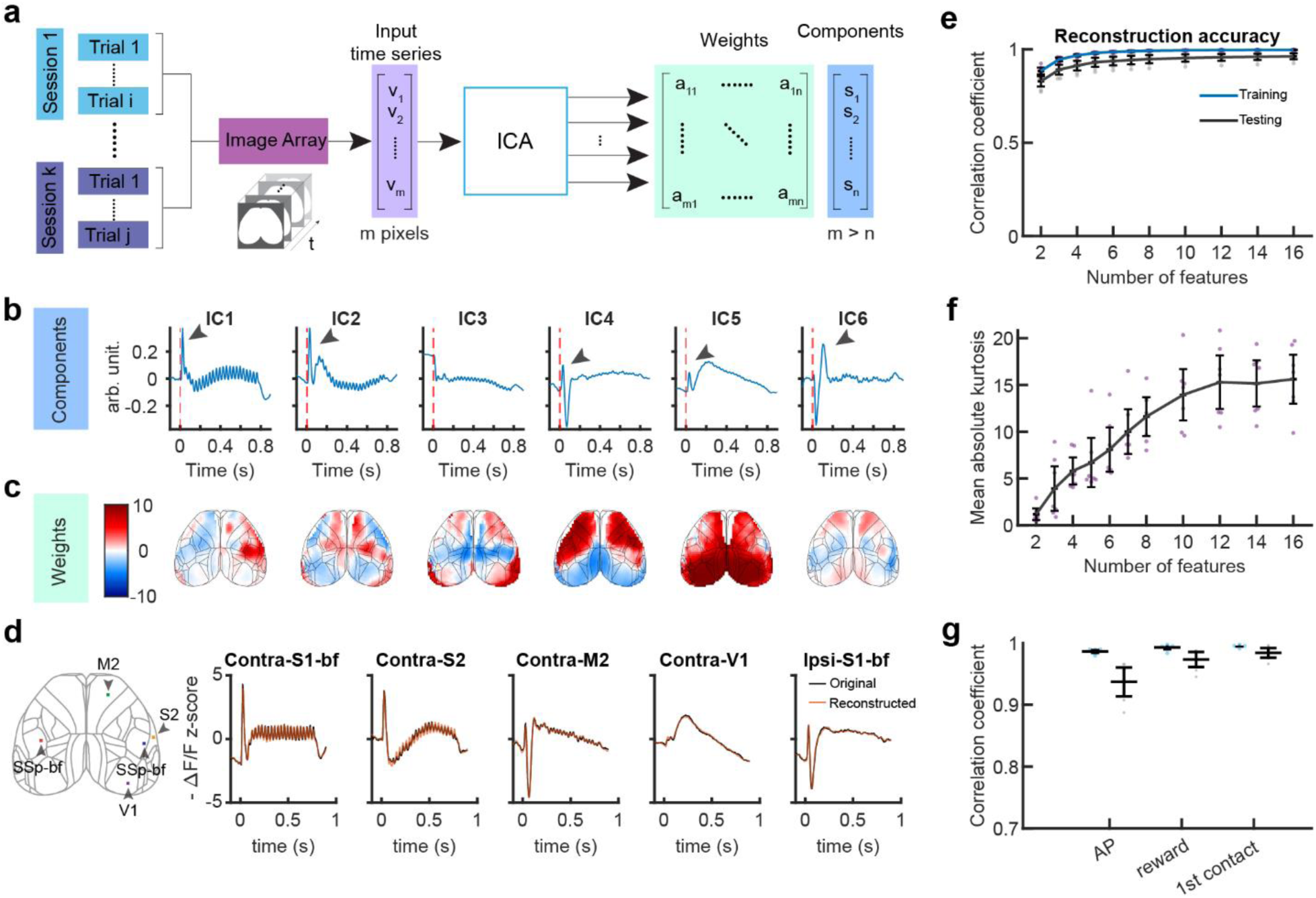
ICA extracts spatiotemporal patterns related to sensory response. (a) Schematics demonstrating how ICA was computed based on average peri-event activity. The ICA is applied on the temporal domain, reducing dimension from around 1400 pixels to 6 components. Non-cortical pixels were masked and not included in the analysis. (b) An example set of temporal ICs during 30 Hz air-puff stimulation. Each IC was normalized by its L2 norm. N = 1 animal, input data were average from 946 trials. (c) Spatial distribution of the weights for each IC shown in (b). (d) Reconstruction of signals using 6 ICs overlapped greatly with the original time-series at the same ROI. The reconstruction was calculated through the linear combination of ICs and their weights shown in (c) and (d). (e) Determination of the number of ICs based on the reconstruction quality. The reconstruction quality is quantified using the correlation coefficient between the original and reconstructed time series aligned with air-puff onset. Trials were randomly shuffled into training and testing datasets iteratively. ICs established from the training dataset were used to calculate the weights of each IC in the testing dataset. Error bar denotes 95% confidence interval (CI). N = 7 mice, where each point represents average correlation coefficients of 10 iterations of an animal. (f) Determination of the number of ICs based on the mean absolute kurtosis. (g) Correlation coefficient between the measured signals aligned to AP, reward and 1^st^ contact and those reconstructed from 6 ICs, for trials in the training (left, blue) and test sets (right, black). The ICA works well for other peri-event alignment with a reconstruction accuracy above 95%.

The corresponding weights of the normalized ICs demonstrated the spatial distribution of each component (**Fig. 4c**). IC1 captured the onset as well as the 30Hz oscillatory responses to the sensory stimuli, and it was most prominent in the contralateral barrel cortex (SSp-bf) as expected. However, a distinct path of contralateral M2 also exhibited a high-frequency response to the air-puff stimuli, indicating a specific functional connection between these areas. IC2 to IC5 showed temporal response profiles to the stimulus onset with various delays, and their spatial locations in frontal, lateral, or posterior parts of the cortex suggested that these networks are engaged in higher-level processing of the stimulus in the behavioral context of the task. Overall, the ICs revealed the fast spread of a whisker stimulus across dedicated cortical subnetworks with shared dynamics that likely correspond to sensorimotor integration.

### Reaching is associated with the activation of a transient sensorimotor network as well as premotor ramping activity

Having established the t-ICA method with neural responses to external stimuli, we next investigated whether this approach could identify subnetworks involved in the preparation and execution of reaching behavior.

For this analysis, we extracted six ICs from trial-averaged activity centered to the moment of first contact with the waterspout **(Fig. 5)**. The six identified ICs **(Fig. 5a)** allowed for a reconstruction of the original signal of all pixels at an accuracy > 95% for the 20% of test-trials not used in the IC computation and near 100% for the 80% of trials that were used (**Fig. 4g** ‘1^st^ contact’). The temporal dynamics of the six identified ICs (shown for one sample mouse in **Fig. 5a**) reveal multiple interesting features, most of which could be identified for each mouse (**Supplementary Fig. 2**). A consistent temporal fast dynamic (IC1) consisting of a fast activity peak initiated upon waterspout contact was present across mice (**Fig. 5b, c**). The weights of this dynamic were largest in primary sensory and motor upper limb areas with some differing amount of participation of premotor regions across animals **(Fig. 5b).** When pixel-wise weights of IC1 from left-and right-side trials were averaged respectively, the spatial weight maps showed more presence of IC1 on the contralateral side of the hemisphere with respect to the side of reaches, indicating contralateral predominance of this temporal dynamic (**Fig. 5d, e**). The lateralization of neural activity to the contralateral hemisphere during reaching movements is difficult to detect using standard ΔF/F imaging alone (**Fig. 1d**), as multiple concurrent neural processes likely overlap within the same cortical regions. By applying ICA, we successfully disentangled these distinct processes, revealing the functional lateralization pattern that corresponds to reaching behavior on a single trial basis. This approach demonstrates how computational decomposition methods can enhance our understanding of the distributed neural networks underlying complex motor behaviors.

**Figure 5:**
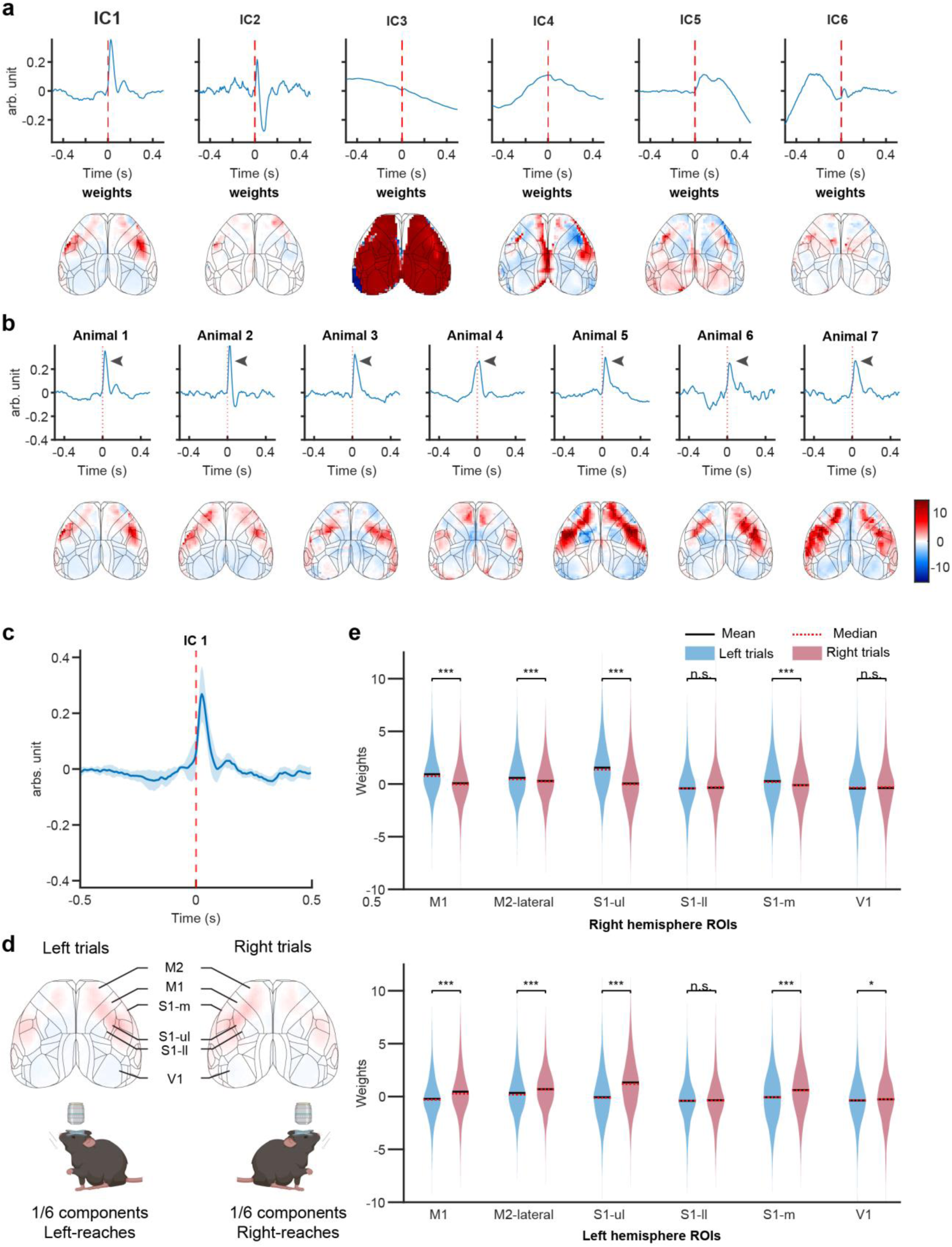
Spatiotemporal patterns capture variabilities in individual trials. (a) A set of six ICs from a representative animal. Top row shows temporal ICs aligned with the first contact from success trials. Bottom row shows the corresponding weight distribution across the cortex. ICA was calculated on mean event aligned data of left and right trials combined for each animal. (b) IC1, a sharp peak in activity upon the first contact with reward, is observed consistently across animals. Top row shows the temporal IC from seven animals, and bottom row shows the corresponding weight distribution across the cortex (c) Average of IC shown in (a) across animals shows consistent alignment with the moment of contact. N = 7 mice. (d-e) The IC is more present on the contralateral S1, M1, and M2 of the reaches. (d) The spatial maps show average weights of the IC across left-and right-reach trials. (e) IC at most ROIs including S1-ul, M1, and M2 on the contralateral hemisphere shows significantly higher weights than that on the ipsilateral hemisphere. N = 3241 left-reach trials and 3179 right-reach trials from 7 mice for both (c) and (d).

### Wide-field voltage activity contains stimulus information throughout task execution and allows prediction of trial outcome

We next examined to what degree the stimulus and behavioral outcome is encoded in the neural activity over the course of the trial. Artificial neural networks have emerged as a method to decode behavioral variables from complex neural activity, with a wide variety of architectures employed^53, 54, 55^. Fully-connected neural networks can be used to test whether nonlinear combinations of input features can be used to test how much information about sensory stimuli or behavioral actions is contained in neural activity^56, 57^. We trained simple feedforward neural networks to perform a binary classification from the neural activity recorded from 36 selected ROIs over 200 ms time windows. For each time window and mouse, a two-layer neural network was trained with stratified k-fold cross-validation (see **Methods**). We examined the accuracy of stimulus classification (left- vs right-reaching) throughout the trial aligned to stimulus presentation, reward, and first/expected contact (**Fig. 6a**). As expected, the accuracy was at chance level before the air-puff stimulus (AP), and highest immediately after. While the accuracy gradually decayed after stimulus presentation, there was a secondary gradual rise that peaked immediately after the response. When classifying the behavioral outcome (success or no-response), the accuracy was significantly above chance throughout the trials, showing that as early as before the AP stimulus, the neural activity can predict the trial outcome with some accuracy (**Fig. 6b**). The accuracy gradually rose until it reached its maximal level shortly after the response.

**Figure 6.**
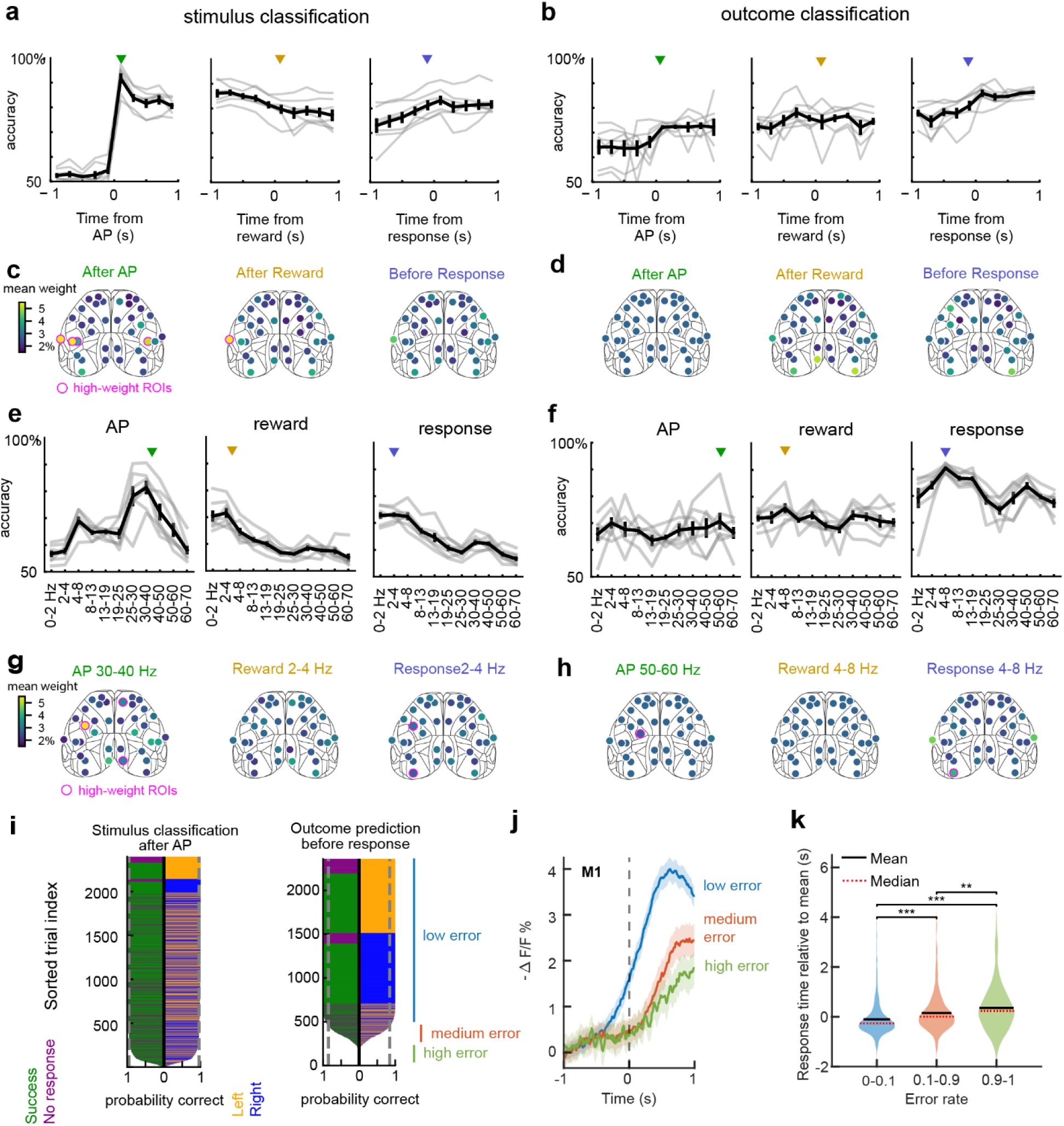
Neural Network predictions from neural activity. (a) Balanced accuracy of neural network classification of stimulus (left or right) from the neural activity of 36 ROIs over non-overlapping 200 ms segments. Gray lines show the trial average of each mouse; black lines and error bars show average and standard error of the mean across mice. Time segments are aligned to AP (left panel), reward presentation (middle), and response (right), where response means either the moment of first contact (for success trials) or 2 s after reward presentation (for no response trials). (b) Same as (a), but for outcome (success or no response) classification. (c) Normalized weights from each ROI after AP (left), after reward (middle) and immediately before response (right). The time segments shown are indicated by the triangles in (a). High-weight ROIs, defined as ROIs with weight distribution significantly different from a shuffled control (P < 10-4), two-sided Mann-Whitney U test) and mean weight at least one s.d. greater than the population mean, are outlined in magenta. See Methods for details. (d) Same as (c), but for outcome classification. Time segments are indicated in (b). (e) Balanced accuracy of network classification of trial outcome (success or no response) from power spectrum. Power spectral data from limited frequency bands across 2 s about each behavioral event was used as input to the classifiers. (f) Same as (e), but for outcome classification. (g) Normalized weights from each ROI for the frequency band with greatest accuracy in each section of the trial. The frequency bands shown are indicated by the triangles in (f). (h) Same a (g), but for outcome classification. (i) Distribution of correct stimulus classification probabilities after AP (left panel) and that of correct outcome prediction probabilities before response (right panel) for example mouse (Animal 1). Each horizontal line represents a trial whose length indicates the probability that it was classified correctly when in the test set (see Methods). The left half of the line is colored according to the trial’s outcome (green if success, purple if no-response) and the right according to its stimulus (orange if left, blue right). The trials are sorted first according to the probability of correct classification, then according to stimulus, then outcome. Dotted line shows average accuracy. (j-k)Trials of low, middle and high error rate for outcome prediction show divergent neural activity and response time. *N* = 1 animal, 2343 trials. See **Supplementary Fig. 6** for all animals. (j) Activation level is higher in trials with lower error rate as demonstrated M1 activity aligned to first contact. Lines denote mean, and shaded areas denote 95% CI. (k) Response time is lower in trials with lower error rate. P values = 10^-18^, 10^-16^, and 0.0032 from left to right, with two-sided Mann-Whitney U Test.

We then examined how much each brain region contributed to the classifications. To do this, we computed the mean absolute-valued outgoing weights from each ROI. The weights were normalized and averaged across multiple repetitions and provide a measure of how much each ROI contributes to classification. By this measure, the networks primarily used information from the sensory cortex immediately after AP and reward to predict the stimulus (**Fig. 6c**). However, in the time window immediately preceding response, the weights became more distributed. We see an opposite trend with outcome classification; weights are initially evenly distributed across ROIs, but became concentrated to retrosplenial, primary visual, and primary motor areas as each trial progressed. However, ROI weights for outcome classification remained evenly distributed throughout the trial compared to stimulus classification: high-weight ROIs, defined as ROIs significantly greater than the shuffled mean (magenta, **Figs. 6c-d**), were present in the stimulus classifiers in the AP and reward periods but were not present in outcome classifiers. This suggests that information on task engagement is distributed across brain regions; this was further supported by classifiers trained on activity limited to left-right ROI pairs, which could match the 36-ROI classifiers only when classifying stimulus from barrel cortex (**Supplementary Fig. 3**)

Next, we examined how information about stimulus and outcome is encoded across frequency bands. For each segment of the trial, the power spectral time course of each frequency band over the 2 second interval was used to train and test neural networks for classification. Accuracy for stimulus classification in the AP interval was greater for higher frequencies and was highest in the 30-40 Hz range. In the reward and response periods, the lower frequency bands were more informative about stimulus, peaking at 2-4 Hz (**Fig. 6e**). For outcome classification, accuracy was similar across frequencies during AP and reward but peaked at 4-8 Hz during the response (**Fig. 6f**). We examined the weight distribution for the frequency band with highest accuracy in each case. Weights were concentrated in the sensory and retrosplenial regions for 30-40 Hz during AP. During the response and reward, information in the 2-4 Hz frequencies were distributed across the cortex (**Fig. 6g**). When classifying outcome, weights are evenly distributed across areas during AP using 50-60 Hz and reward using 4-8 Hz. During reward using 4-8 Hz, weights become concentrated to the visual and motor regions (**Fig. 6h**)

We then asked whether there were trial-by-trial differences in classification accuracy: did all trials have a similar probability of being classified to the correct stimulus/outcome, or were some trials easy to classify and others difficult? To answer this question, we examined the probability that each trial was correctly classified in a test set prediction. Specifically, we were interested in examining the distribution of correct classification probabilities across trials and whether they were concentrated around the mean value, or whether the distribution revealed high- and low-accuracy trial groups. We found that when classifying stimulus from activity immediately after AP, most trials have a higher-than-average probability of correct classification, and a small set of trials have very low probability (**Fig. 6i**, left). This trend was also seen when predicting outcome from activity before the response (**Fig. 6i**, right). We can thus divide the set of trials into groups with low, medium, and high error rates (shown in cyan, red and green bars respectively in **Fig. 6j**). Similar distributions were seen across all mice (**Supplementary Figs. 4, 5**).

Trials in the low error rate group in the above analysis were consistently classified correctly across all train-test splits while those in the high error rate group were consistently classified incorrectly, with those in the medium group in between. We examined whether there were any differences in neural activity or behavior across these sets of trials (**Fig. 6j, k and Supplementary Fig. 6** for individual mice). We assessed activation in the primary motor cortex of successful trials within each trial group aligned to first contact (**Fig. 6j**). The low error-rate trials displayed a characteristic ramping-up in activity preceding the contact and had an overall higher activation than the medium and high error rate groups. Furthermore, there was a significant difference between the response times of each trial group: the low error-rate trials had the fastest responses, and the high error-rate trials had the slowest (**Fig. 6k**). This shows that most successful trials are characterized by a ramping up in the motor cortex before contact and fast response times; the neural network correctly classified these trials with very high accuracy. On the other hand, there were a set of successful trials in which the ramping was absent, response times were slower and were consistently classified as no response by the neural networks. This ramping of activity in M1 was a widespread, but not universal, feature used by the classifiers in outcome prediction.

## Discussion

We imaged pan-cortical activity of layer 2/3 pyramidal neurons with the JEDI-1P voltage sensor allowing unprecedented monitoring of high-frequency and subthreshold network activity. We were able to identify gamma oscillations behaviorally locked to reward presentation in frontal premotor areas and a large-amplitude 8Hz oscillation linked to a behaviorally disengaged state with pronounced lack of movement. Other networks were activated with non-oscillatory dynamics, and a given cortical area could take part in multiple such networks identified by their shared temporal dynamics. These dynamics often included fast components such as stimulus locked oscillations and transient activations linked to stimuli or movement. Many such networks showed bilateral activation of the same functional areas indicating widespread bilateral fast dynamics in rodent task processing.

Synchrony in distinct frequency bands (aka ‘oscillatory network activity’) has long been proposed to be involved in information processing and communications between brain regions^2, 58, 59, 60^. In our study, the increase in gamma power at M2 during success trials but not no-response trials indicated its potential role in attention or sensorimotor processing. The source of cortical gamma oscillations has been attributed to fast-spiking PV interneurons and its inhibitory connections with L2/3 and L5 excitatory neurons^11, 13, 61, 62^. Existing cell-type specific widefield calcium imaging of GAD2 neurons of a similar goal-directed task has also shown activation patterns of inhibitory neurons bilaterally in M2 during the response period^23^. Given inhibitory-excitatory dynamics driven by fast-spiking PV interneurons, the activation of inhibitory neurons specifically at frontal M2 could explain the gamma oscillations we observed in relation to successful reaches. The persistent timing of observed oscillations after reward consumption suggests an involvement in updating prior trial information that is widespread in motor and frontal areas^63^ after the completion of each trial. The gamma rhythm, specifically, has been hypothesized to support mechanisms of plasticity, as gamma cycles may generate spike-time dependent plasticity windows^64^ and could result in long lasting task expectations.

Voltage imaging revealed a strong ∼8Hz oscillation in frontal sensorimotor areas that was strongly associated with an akinetic motor state (**Fig. 3**). The persistence of this oscillatory state beyond the go-cue was correlated with reach movement failure. As such this state resembles the features of a disengaged task performance state that has been previously identified with GLM-HMM models as a distinct behavioral state^44^ and cortical activity imaged with GCaMP6 during a reaching task revealed higher variability during disengaged trials in sensorimotor cortex^65^. While calcium imaging cannot reveal single trial oscillatory activity, these findings map well onto our results. The presence of a distinct ∼8Hz oscillatory state links task disengagement to previous human literature employing MEG and EEG methods that revealed an alpha-mu (7-14 Hz) rhythm linked to a lack of attention^66^, localizes to sensorimotor cortex and desynchronizes during movement processing^17, 67, 68^. Thus, voltage imaging enables the mechanistic study of task-related oscillatory activity observed in human MEG and EEG studies with the modern mouse toolkit of causal brain manipulations which will allow the exploration of the mechanistic basis and causal influence of brain oscillations going forward.

Our study revealed distinct temporal dynamics for specific cortical subnetworks related to executing a sensory cued reaching task using temporal ICA. When network activity was aligned to specific behavioral events, multiple subnetworks each identified by their own dynamics related to the behavioral event existed in parallel, and individual brain regions could participate in several subnetworks. One observed coding principle was the predominant bilaterally symmetric activation in each task related network, even when the applied stimulus or the executed movement were unilateral. Bilateral symmetry is commonly observed with wide-field calcium imaging^24, 40^, and in default mode or task related cortical networks identified by fMRI methods^69, 70, 71^ as well as in dynamic patterns found in fMRI signals^72, 73^. Our previous work showed that at a slower time scale, hemodynamically defined bilateral networks are well aligned with voltage imaging using a Butterfly 2.1 sensor^74^. Voltage imaging with the faster JEDI-1P sensor allowed us to determine that the temporal precision of such bilateral activation far exceeded the resolution allowed by fMRI or calcium imaging that also observe such bilateral activity patterns in similar goal-directed tasks. Notably, we found that even those networks showing fast and precise temporal dynamics connected to the reaching movement comprise sensory and motor cortical functional areas. Thus, sensory and motor processing was integrated at all time scales. It should be noted that proprioceptive information is coded both in S1 and M1 in mice^75^, and our fast dynamic reaching networks may well be related to proprioception.

We also found t-ICA networks with slower activation dynamics preceding reach onset (**Supplementary Fig. 2**, IC4). This network dynamic had clear participation of M2 in most mice, with more variable contribution of other areas. Previous work has indicated that such motor preparatory activity builds up across cortical areas during learning and is organized by M2 inputs^27, 76^. In agreement with these studies, our behavioral response prediction using neural networks trained on voltage imaging data just before the response (**Supplementary Fig. 3**) showed that M2 activity retained the most information about stimulus classification at this time while also predicting response outcome. However, the distributed nature of information processing was also clearly apparent in the neural network prediction analysis, as the activity from many cortical areas could classify the stimulus as well as the behavioral outcome well above chance.

Overall, our results revealed multiplexed cortical functional networks with subnetworks sharing specific and sometimes oscillatory dynamics. As JEDI-1P primarily reveals subthreshold membrane voltage dynamics and therefore synaptic activity, the signal shares dynamics with the local field potential^39^. Due to the Kv2 tag of the vector, our imaging was specific to somatic voltage changes in L2/3 pyramidal neurons and revealed fast dynamic synchronous synaptic activity of these neurons specifically. Key messages of behavioral control we found are that gamma oscillatory dynamics may be specific to rule updating in frontal cortex, that prominent slow ∼8 Hz mu-oscillations shared between somatosensory and motor areas relate to the disengaged state, and that sensorimotor integration between areas is synchronized at least down to the ∼10ms time scale that we could resolve. Likely the networks we revealed are coupled not just through cortico-cortical connections, but also subcortical loops. Future work with simultaneous deep recordings will be needed to unravel such vertical organization.

## Methods

### Animals

A total of 7 EMX1-Cre mice (5 males and 2 females) was used for this study. Mice were maintained at 22 °C with the humidity ranges from 30–70%. Post head-plate implantation surgery, animals were singly house with enrichment under a reverse light cycle (12 h light, 12 h dark). Training and imaging experiments were performed during the dark cycle. Animals were provided with ad libitum food and water prior to the surgery and behavioral training. After surgery recovery, animals were placed on a continuous water-restriction schedule. Emory University Institutional Animal Care and Use Committee approved all animal work performed in this study.

### Neonatal intracerebroventricular injection

For widespread expression of JEDI-1P. and reference fluorescence throughout the brain, intracerebroventricular injection was employed to administer viral vectors^77^. Breeding pairs were established using EMX1-Cre mice and monitored breeder cages twice daily as the expected delivery date approached. Upon observing visible milk spots, we carefully removed newborn pups from the breeder cage and placed them on a 37 °C heating pat during viral vector preparation. The viral mixture was consisted of AAV.PHP.eB-EF1a-DIO-JEDI-1P-Kv2.1-WPRE (4 × 10^12^vg/mL) and AAV9-hSyn-mCherry (1.3 × 10^13^ vg/mL) at a 1:1.3 particle ratio. For each pup, 5 µL of the viral mixture was loaded into a 10-µL Nanofil syringe (World Precision Instrument). The injection site was identified as 2/5 of the distance between the lambda suture and each eye. The syringe was slowly and steadily inserted to a depth of 3mm at the targeted site, and 2 µL of viral vectors were administered into each hemisphere. Immediately following completion of all injections, pups were returned to the breeder cage to be cared for by the breeders.

### Surgery of imaging window and head-post implant

The imaging window was adapted from the clear-skull techniques described in existing imaging studies^39, 78, 79^. Anesthesia was induced using 3% isoflurane and maintained at 2% throughout the surgical procedure. Nair (I0041395, Church & Dwight) was used for fur removal over the scalp and the skin around the neck. 70% ethanol and povidone-iodine pads were alternated a total of three times ending with povidone-iodine to clean the scalp. The scalp was then carefully removed to uncover 9 × 9 mm of the skull. To increase the adhesion of the dental cement, the fascia over the skull was gently scraped off with a surgical scalpel, and the skull was lightly scored with the scalpel and then dried with cotton swabs. A thin layer of Opti-bond Universal (36519, Kerr) was applied to the skull surface and then cured with ultraviolet light. A different mixture of dental cement was then applied over the cured the Opti-bond surface. The new mixture was consisted of 1 scoop of C&B Metabond L-powder (S399, Parkell Products Inc.) and 5 drops of C&B Metabond Quick Base (S398, Parkell Products Inc.). A custom cut coverslip (12-545-88, Fisherbrand Superslip) with a shape of hexagon was quickly placed over the dental cement while it was still in its liquid form to allow adjustment of the glass placement if necessary. The gap between the coverslip and the skull was carefully filled with rest of the dental cement to avoid any air bubbles. After the dental cement under the imaging window was sufficient cured, the custom designed headplate for head-fixation was placed around the imaging window and secured with an opaque mixture of the dental cement (2 scoops of L-powder and 6 drops of Quick Base). Following the surgery, animals were placed in a clean cage over a heating pad at 37°C and were monitored closely until they were fully awake and mobile.

### Continuous water restriction

Mice underwent continuous water restrictions at least 2 days before the experiments to ensure a steady weight. On the first day of water restriction, the ad libitum water was removed. At the same time, the body weight was measured and used as the baseline weight for future reference. The animal was weighed daily and provided with a minimum of 40 ml/kg hydrogel water, calculated using the baseline, to maintain above 80% the baseline weight.

### Bilateral reaching task and behavioral training

The reaching task was designed to assess neural activity of movements for both forelimbs. Animals were trained to reach for sucrose water with the limb on the same side as the reward. A typical trial in a fully trained session consisted of baseline, air-puff train stimulation of either 30 Hz or 50 Hz, delay period with a variable length, visual cue, reward dispensation, response period, and inter trial interval (ITI). In all mice 30 Hz air-puff stimuli were applied on the left whiskers, and 50 Hz air-puff stimuli were applied on the right whiskers. In 4 out of 7 mice the left whisker cue was followed after the delay by a left-side water reward, and the right-side cue by a right-side reward. These 4 animals had also previously undergone training to lick the same-side tube to initiate the reward and obtain it. In the remaining 3 mice this cue and reward sides were reversed, and previous lick-training did not take place. A success trial was defined as a trial where contact with the waterspout where the reward was present. A trial was labeled as no-response failure if the reward was delivered but not received. A trial was considered early-contact failure if the animal touched the waterspouts before end of delay period. A total of 6777 success trials and 1629 trials were used in analyses. Early-contact failures were not discussed in this study.

To reduce the level of stress of the animals, the mice were habituated to the handling and head-fixation for up to five days. To first form the association of the air-puff stimulation and the side of the reward, the two waterspouts were first placed close to each other and were placed within licking distance of the animals to encourage reward consumption. After animals were able to receive reward reliably on both sides with their tongue, the distance between the waterspouts increased incrementally and further away from the animals’ mouth. There was an initial drop in performance before the animals were accustomed to reaching for the reward with their limbs. If the animals did not show any sign of reaching, gentle air was applied on the whisker pad on either side to evoke grooming which subsequently led to reaching.

Note that a 0.5 s period following the go cue was originally designed to encourage the animals to perform a decision reach to trigger the reward. If the subject touches the waterspout on the same side as the air-puff stimuli, reward was immediately dispensed.

However, none of the animals learned to perform this action with high enough performance accuracy. Only 1.9 % of the success trials, 167/8637 trials, were triggered by contact during the 0.5 s visual cue period. These trials were removed from analyses to keep the experimental condition consistent.

### In vivo wide-field imaging

Wide-field imaging was conducted at 200 Hz using a commercial CMOS high-speed system (MiCAM ULTIMA. SciMedia Ltd) with dual cameras (**Fig. 1a**). Each imaging trial was 10.24 s in duration. Both JEDI-1P and mCherry fluorescent protein were excited with a single blue LED light (466/40 nm, center wavelength/bandwidth FF01-466/40, Semrock), through the intact skull and the imaging window. The excitation light power at the plane of the imaging window 0.05 mW/ mm2 for all imaging sessions. The excitation light was reflected using a dichroic beam splitter at 495 nm (FF495-Di03-50×70, Semrock). The emission light from JEDI-1P and mCherry was split into two using another dichroic at 580 nm (FF580-FDi01-50 × 70, Semrock). The split emitted light from JEDI-1P and mCherry was conditioned by 525/50 nm (Semrock: FF03-525/50-50) and 650/60 nm (FF01-650/60-50, Semrock) emission filters, respectively. The field of view of each imaging camera was 10 × 10 mm in size with resolution of 100 × 100 pixels (0.1 mm/pixel). 3D printed light barrier along with pliable putty (10-2625, TheraPutty) were used to shield ambient light from contaminating the imaging. The barrier also functioned to minimize the influence of blue excitation LED on the pupil size and the behaviors.

### Design of behavioral setups

Behavioral setups were custom built to accommodate the bilateral reaching task with synchronous neural recording and behavioral tracking. Each behavioral rig was consisted of four main components: head-fixation station, sensory and reward delivery system, contact tracking sensors, and video tracking with multiple cameras.

The head-fixation station was designed to minimize motion artifacts. Animals were situated in a 3D printed tube tilted at 45° and were presented with resting bar in front to improve comfort and reduce stress during the behavioral sessions. A custom-made holder was used to secure two tubes that delivered air-puff train and LED light panel that omit visual cues. The gentle air puff was achieved by driving a miniaturized solenoid valve (LHDA1233215H, The Lee Company) with TTL pulses at 30 or 50 Hz to release pressured air. For sugar reward delivery, two waterspouts were connected to a gravity-based reservoir and were each controlled by the same type of solenoid (LHDA1233215H, The Lee Company) for small volume water dispensation. To adjust the waterspout distance at ease during training, the two waterspouts were attached to a pair of scissor-like arms. A capacitive touch sensor was connected to the metal part of each waterspout to register the contacts between the animal and the waterspouts. Two additional touch sensors were connected to the resting bars to track contact with the bars. Lastly, multiple cameras (Model numbers, Basler) were configured to capture animals’ facial and limb movements from different angles.

### Synchronization across devices

At the center, Bpod (Bpod State Machine r2, Sanworks) functioned as the primary device that synchronized all components. Aside from controlling the state flow of the designed task, Bpod sent digital signals to trigger external devices. Digital TTL pulses were used to control visual cues, the opening and closure of solenoid valves, and frame by frame captures of the cameras. Additional, Bpod received signals from external devices to achieve cross-device synchronization. The timestamps of both the input and output signals were used to align data with different modalities to the same timeline.

### Behavioral recording

All imaging sessions were synced with at least two cameras (a2A1920-160um, Basler and M3Z1228C-MP, Computar) to record the animal behavior. One camera faced the front left of the animal, and the other one captured the bottom view of the mouth at about a forty-five-degree angle facing upward (**Fig. 1c**).6527 trials from 4 out of 7 animals also contained recording of the left eye to track pupil diameter. IR LEDs (LED780E and LEDMT1E, Thorlabs Inc) were used to provide sufficient light for the behavioral cameras without interfering with the voltage imaging.

Behavioral videos were acquired at 50 FPS or 100 FPS and saved in .mp4 format. The TTL pulses that used to trigger image capture were stored in Bpod and were used to splice videos into individual trials aligned with the onset and offset of the imaging data.

## Co-registration of imaging data across sessions and subject

Running statistical analyses reliably across sessions and across animals requires registering all imaging data to the same standard cortical map outlined from the Allen Mouse Brain Common Coordinate Framework^80^. Subtle x and y axis shift might also happen between the two CMOS cameras over time. To better align imaging data across trials, the co-registration was achieved semi-automatically through four steps. First, average frames plotting ΔF/F in response to air-puff onset from an example session per animal were calculated and output as 50 × 50-pixel images. In Step 2, the activation at SSP-bf on either hemisphere were used to shift these images to the standard cortical map manually to help generate the template image for each animal. The template images were represented using raw image of the cortex from the JEDI-1P channel which best captures the location of major blood vessels. In Step 3, 2D cross-correlation was run between the JEDI-1P and the mCherry frames from the same trial to calculate the x and y offset. The offset was then used to align the JEDI-1P images with that of the mCherry frames. The cross correlation was calculated independently for five randomly selected trials per session. The modal offset for each axis to minimize errors. In Step 4, 2D cross-correlation was calculated between the template and the shifted JEDI-1P frames for each imaging session. The modal offset between the template and the JEDI-1P channel within each session was used to align session-wide data to the same map automatically.

### Preprocessing of imaging data

The preprocessing pipeline was developed to remove hemodynamic, motion, and photobleaching artifact by utilizing the reference channel to subtract non-voltage signals from JEDI-1P traces^39^. The co-registration data were used to first align all imaging frames to the same standard map. Raw imaging signals from both channels represented the absolute light intensity of the omitted light. The signals were first subtracted by background fluorescence, 590 arbs. unit for JEDI-1P and 270 arbs. unit for mCherry, measured from control animals without fluorescent expression. Photobleaching was subtracted from the JEDI-1P channel using an exponential decay model that estimates the photobleaching over 10.24 s. Then (F – F0)/F0, often known as ΔF/F, of both channels was calculated where the baseline F0 was defined as the average intensity from 1.2 to 1.7 s to avoid fast photobleaching during the first 1 s of the imaging. The reference ΔF/F was filtered at different frequency bands to extract non-voltage signals at [10, 30], [1, 10], and below 1 Hz. The filtered reference traces were then used to regress artifacts at each frequency range from the JEDI-1P traces sequentially using the ordinary least square methods. The preprocessing was performed at binned pixels with a size of 2 × 2 pixels accelerated with GPU. The resulting ΔF/F traces have been demonstrated to effectively remove most artifacts and reveal relevant voltage signals in our previous study.

### Peri-event activity alignment

Given that there was variability in delay period duration and response time of reaching, voltage traces were aligned with specific events with a fixed duration before and after the aligned time point for further analyses. Specifically, two-second activity was centered around the air-puff onset where the second before the onset was used to represent baseline and the second following the onset represent the stimulus duration. Two-second activity was centered around the reward dispensation for both success and no-response trials. For the response period, the event was centered to the moment of first reward contact in success trials. As there were no valid contact with reward in no-response trials, 1 to 3 seconds following the reward dispensation was used to evaluate activity during response period in these failure trials. This alignment for failure trials was guided by the median time, 1.2 s, it took for all animals to touch the waterspout after the reward delivery.

### Power analysis of peri-event imaging data

Short-time Fourier Transform (STFT) was used to assess the power changes at different frequency ranges by window sizes for each frequency band. The frequency ranges include theta, alpha, beta, and gamma bands. In addition, the power at 6-10 Hz was also assessed to focus on the large-amplitude oscillations at the range. Beta and gamma ranges were further divided into several bands to increase the resolution in the frequency space. Specifically, the following frequency bands and their associated window size were used: theta: 4-8 Hz (0.64 s window); alpha: 8-13 Hz (0.64 s window); 6-10 Hz (0.64 s window); beta: a. 13-19 Hz (0.32 s window) b. 19-25 Hz (0.32 s window) c. 25-30 Hz (0.32 s window); gamma: a. 30-40 Hz (0.16 s window) b. 40-50 Hz (0.16 s window) c. 50-60 Hz (0.16 s window) d. 60-70 Hz (0.16 s window). The overlap between window was 50 %. For a given frequency range, MATLAB function *spectrogram()* was used to calculate the power changes with the associated window size. Then function *bandpower()* was used to calculate the average power, P, within the frequency range. The average power was then converted to decibel scale using 10*log10.

The power analysis was applied on peri-event activity of masked pixels on trial basis. Average power of 1 s before the air-puff onset was used as the baseline. Time series of power during each event were subtracted by the baseline power of the corresponding frequency range at the same pixel from the same trial. The power differences were used to show changes in power in **Fig. 1e and 2f**.

To calculate power spectral density (PSD), MATLAB DSP System Toolbox (dsp) was used Specifically, we used dsp.SpectrumAnalyzer() with Welch method and a window length of 400 samples. The PSD was calculated for individual trials in a batch of 100.

### Large-amplitude oscillation detection

6–10 Hz oscillations that showed large amplitude were detected using a combination of criteria. First, event-aligned ΔF/F traces of bilateral M2 went through a lowpass filter of 10 Hz to remove high frequency activity. Maxima and minima were identified using *islocalmax()* and *islocalmin()*, respectively. Oscillations were considered detected in trials when two criteria were met in M2 regions of both hemisphere: (1) the ΔF/F amplitude between paired peaks and troughs exceeded 4%, and (2) the time interval between consecutive peaks and troughs matched the expected 6–10 Hz oscillation period Alternatively, if the power between 6–10 Hz at bilateral M2 was both above –2dB and the ΔF/F difference exceeded the same threshold, the 6–10 Hz oscillations were also considered to have occurred in those trials.

### Independent component analysis of peri-event imaging data

As shown in **Fig. 3a**, t-ICA analysis was performed on average peri-event activity per subject. For each animal, the average peri-event activity across trials within the same experimental condition was first masked to remove non-cortical pixels, resulting in *m* time series. ICA, *pica()*, was applied on the temporal domain of the matrix to reduce the dimensions from *m* to *n* pixels, where *n* was significantly smaller than the start dimension *m*. Each IC was scaled to unit length with respect to the L2 norm. The weight matrix was scaled accordingly based on the scaling factor of each IC. To determine the appropriate number of ICs, trials were randomly divided into 80 % training dataset and 20 % testing dataset. For each iteration, the set of ICs were used to estimate the weight of each component in the testing data using linear regression *regression()*. The pair of ICs and weights in both training and testing dataset were used to reconstruct pixel-wise signal. Pixel-wise correlation coefficient between the original and the reconstructed signals was calculated to quantify the reconstruction quality of each dataset using *corrcoeff()*. The resulting correlation coefficients for training and testing datasets across ten iterations within an animal were averaged to show how well ICs represent majority of the data. To make the ICs minimally gaussian collectively, the mean absolute kurtosis of the ICs was calculated using *kurtosis (),* where the output kurtosis value was subtracted by 3.

### Quantification of behavioral data

Both limb positions and pupil diameters were estimated with DeepLabCut 2.2 (DLC). The bilateral limb positions were recorded with the bottom-view camera. Multiple points on each limb included the digits, center of the paw, wrist, and points that outline the outer shape of the paw. The estimated points output from DLC were then processed in MATLAB. Instead of relying on the accuracy of individual points, a voting mechanism using median values of all available points was used to approximate the center of the paw position. The estimated paw center was used to calculate the speed of each limb. The histogram of limb speed was used to help identify filter out speed values that are abnormal.

The left pupil was tracked with the side-view camera. DLC labeled 8 points round the pupil. The labeled points with above 0.6 confidence level were used to fit a circle and estimate the pupil diameter. Frames where fewer than 5 points were above the set confidence level were omitted to avoid erroneous pupil diameter estimation.

### Neural network classifications from neural data

Feedforward neural networks were trained to perform binary classification of either stimulus (left or right) or trial outcome (success or no-response). The input was either a segment of the ΔF/F time series or power spectrum data P, both derived from event-aligned voltage traces from 36 selected ROIs.

When using the ΔF/F time series, the signal was first preprocessed by taking the z-score with respect to the distribution of the signal during a pre-trial resting period. The input layer had 1440 units (40 time points from 36 ROIs). The sampling windows were non-overlapping sections of the time series data aligned to a behavioral time point (t = 0). For the air-puff-aligned (AP) windows, t = 0 indicates the moment of AP presentation. For the response-aligned windows, t = 0 indicated either the moment of the mouse’s first contact with the lever for successful trials, or 2 s after the end of the delay period for no-response trials. Ten non-overlapping windows were used with t = 0 as the fifth endpoint, i.e., the windows were, in interval notation, [–1 s, –0.8 s], [–0.8 s, –0.6 s], …, [–0.2 s, 0 s], [0 s, 0.2 s], …, [0.8 s, 1 s]. When using the power spectrum, the power signal from 36 ROIs of a single frequency band over the 2 s behaviorally aligned window was given as input. The input layer size thus depended on the frequency band selected. For both input types, the network had two hidden layers whose sizes were optimized via cross validation.

Models were trained for each mouse and time segment or frequency band separately. First, the trials were split into train and validation sets (8:2 ratio). Because the distribution of trial outcomes was imbalanced, a stratified shuffle split was used. The binary cross entropy was optimized using the ADAM algorithm. Hyperparameters were optimized via a grid search. The hyperparameters tested were as follows: sizes 64, 128, 256, and 512 were tested for the first hidden layer; 8, 16, 32, and 64 for the second hidden layer; the interval [10^−4^, 1] for the l2 regularization parameter (5 steps log-evenly spaced); and [10^−6^, 10^−1^] for the learning rate (6 steps log-evenly spaced). The hyperparameters with the highest balanced accuracy (to account for imbalanced outcomes) on the validation set was selected.

After hyperparameter optimization, the full set of trials were split (stratified shuffle) into training and test sets (8:2). A network was initialized and trained with the selected hyperparameters over the training set and tested over the test set. This procedure (model initialization, train, and test) was repeated 20 times. Then the full test procedure, starting from the train-test split, was repeated 200 times. When reporting classification accuracy, we used the average balanced accuracy over 800 (20x200) repeats. We computed the rate of correct classifications (P(correct)) by dividing the number of times the trial was correctly classified in the test set by the total number of times the trial was in the test set (800 on average).

To compute the normalized weights from each ROI, we initialized and trained the model as in the testing procedure. For each ROI, we found the maximum absolute weight value among outgoing connections from the corresponding input layer to the first hidden layer. The weight values were normalized by dividing by sum of (maximum absolute) weights across all ROIs. This procedure was repeated 50 times for each mouse.

Pytorch v. 2.1.0^81^ was used for neural network training and sklearn v. 1.5.1^82^ was used to generate training set splits.

### Statistics and reproducibility

The behavioral training and imaging experiments in were repeated in 7 mice.

## Data Availability

Imaging data along with the behavioral data will be available in the NWB (Neurodata without Borders) format on DANDI upon acceptance.

## Code Availability

The code for preprocess and analyze the data will be available on GitHub upon acceptance. MATLAB code was tested on version R2023b.

## Acknowledgement

Many thanks to Francois St. Pierre and his lab for providing the JEDI-1P constructs. Dr. Ben Dichter and CatalystNeuro helped with the initial conversion scripts for imaging data to the NWB data format.

## Contributions

D.J. and Y.W. conceived and oversaw the project. Y.W., S.H.K., H.C., and D.J. prepared figures and wrote the manuscript. Y.W. designed and built the hardware and software platform for the behavioral experiments. Y.W. collected the in vivo imaging data. Y.W., E.J.H., and B.L.C handled and trained the animals. Y.W. and B.L.C labeled and refined DLC models to track limb position and pupil diameter. Y.W. and S.H.K. developed code for data analyses.

