## Supplemental Figures for "Cortex-wide voltage imaging in a sensory cued reaching task reveals fast subnetwork dynamics"

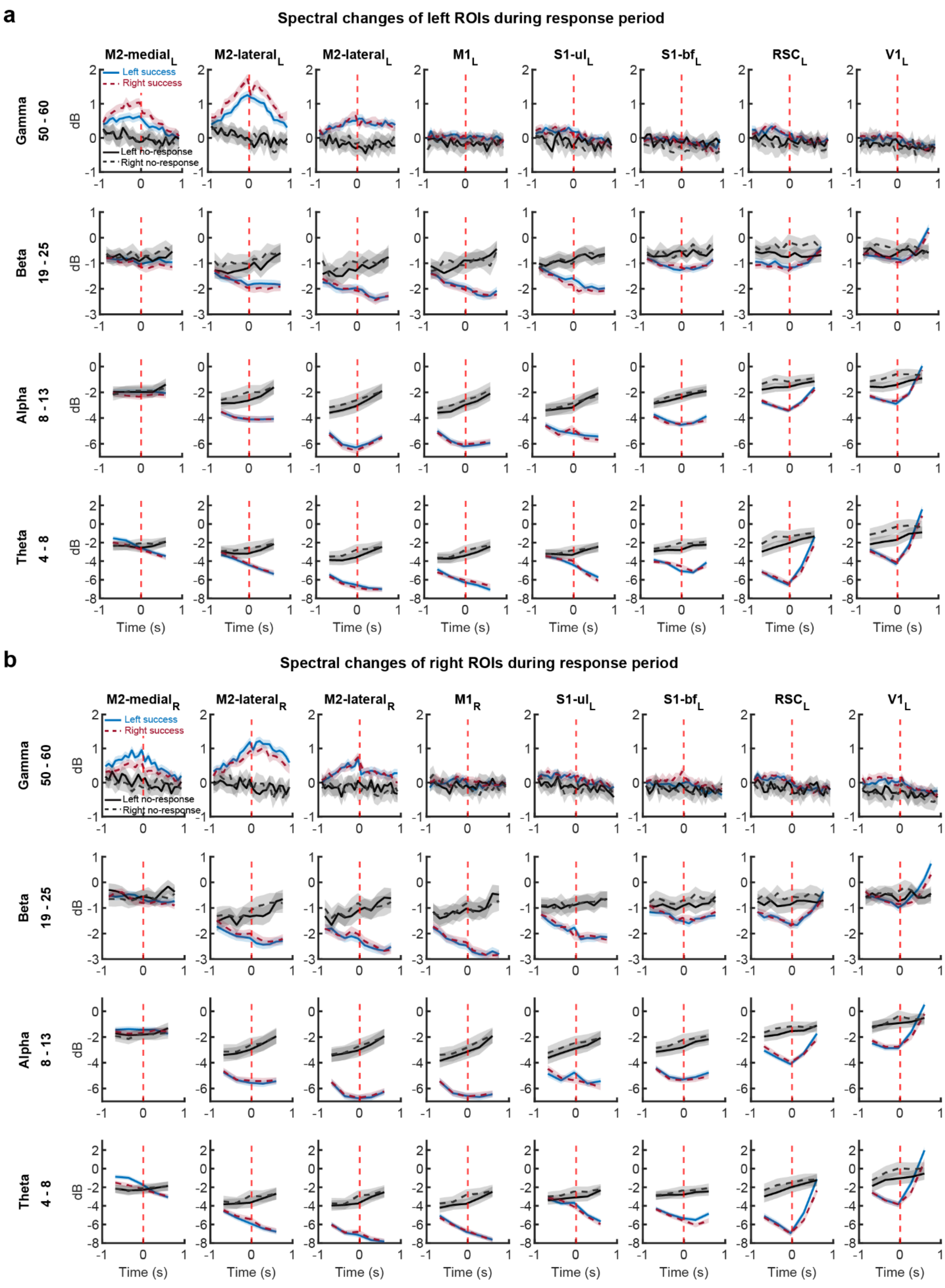
a
Spectral changes of left ROIs during response period
M2-medial L
M2-lateral L
M2-lateral L
M1 L
S1-ul L
S1-bf L
RSC L
V1 L
Left success
Right success
Left no-response
Right no-response
Gamma 50 - 60
Beta 19 - 25
Alpha 8 - 13
Theta 4 - 8
dB
Time (s)
b
Spectral changes of right ROIs during response period
M2-medial R
M2-lateral R
M2-lateral R
M1 R
S1-ul L
S1-bf L
RSC L
V1 L

**Supplementary Figure 1.** Changes of power at different frequency ranges are bilateral with subtle lateralized differences.

(a) Contralateral M2 shows slightly higher gamma power increase in success trials during the response period. Power spectral changes at other frequency bands are mostly bilateral with little differences between left and right trials. Success trials have lower power at these lower frequencies than no-response trials. Red dashed lines are aligned with the reward delivery. ROIs are all from left hemisphere. Solid and dash lines indicate mean power changes at each frequency range. Shaded areas denote 95% CI. N = 3241, 876, 3179, and 753 trials for left-side success trials, left-side no-response trials, right-side success trials, and right-side no-response trials, respectively.

(b) Same as (a) but showcases equivalent ROIs from the right hemisphere from the same set of trials.

First contact tICA of individual animals - left-side success trials

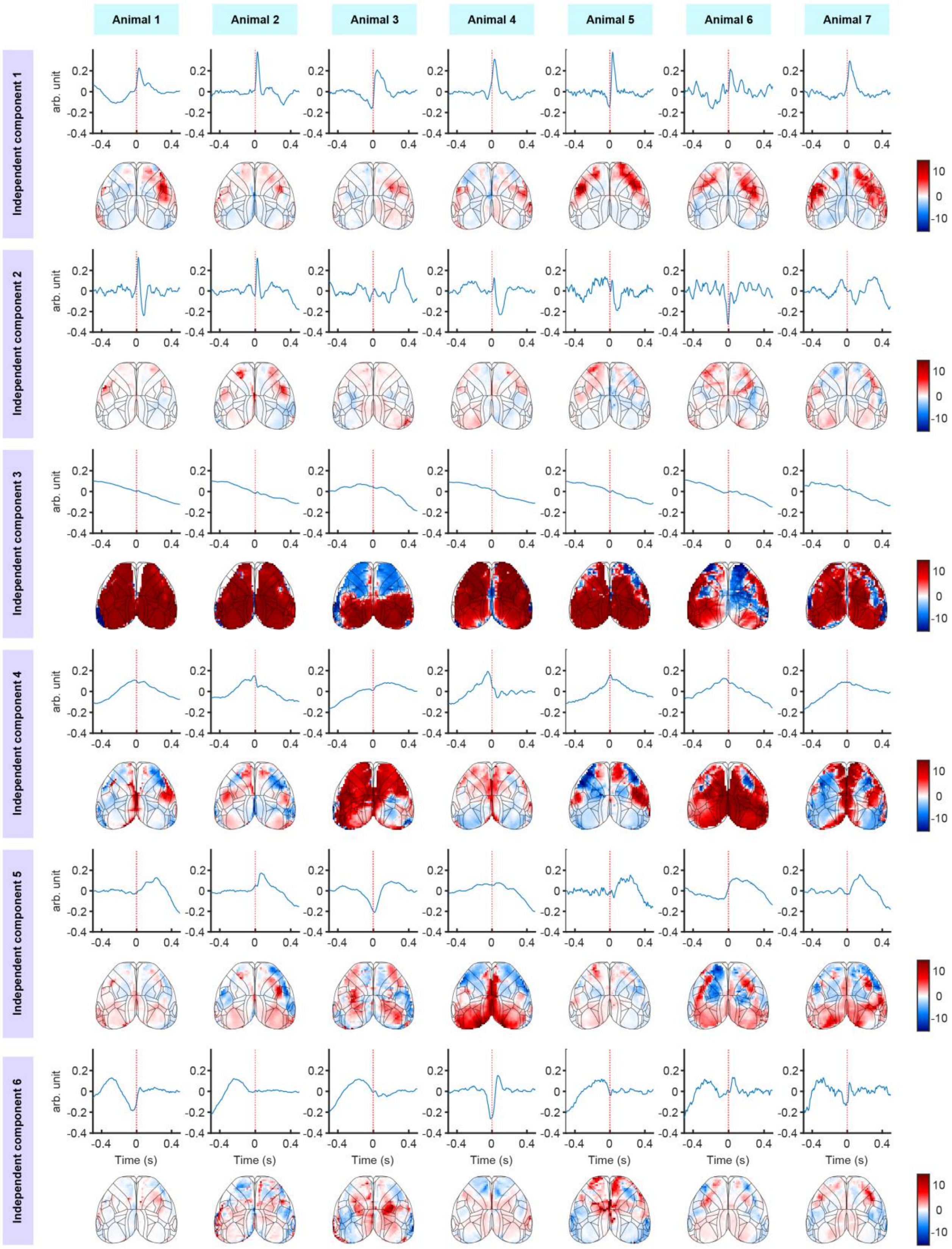

**Supplementary Figure 2.** ICA aligned with response period from all 7 mice showed similar spatial temporal patterns. Each row represents one of the six ICs. Each column shows the extracted IC and its corresponding spatial map per animal.

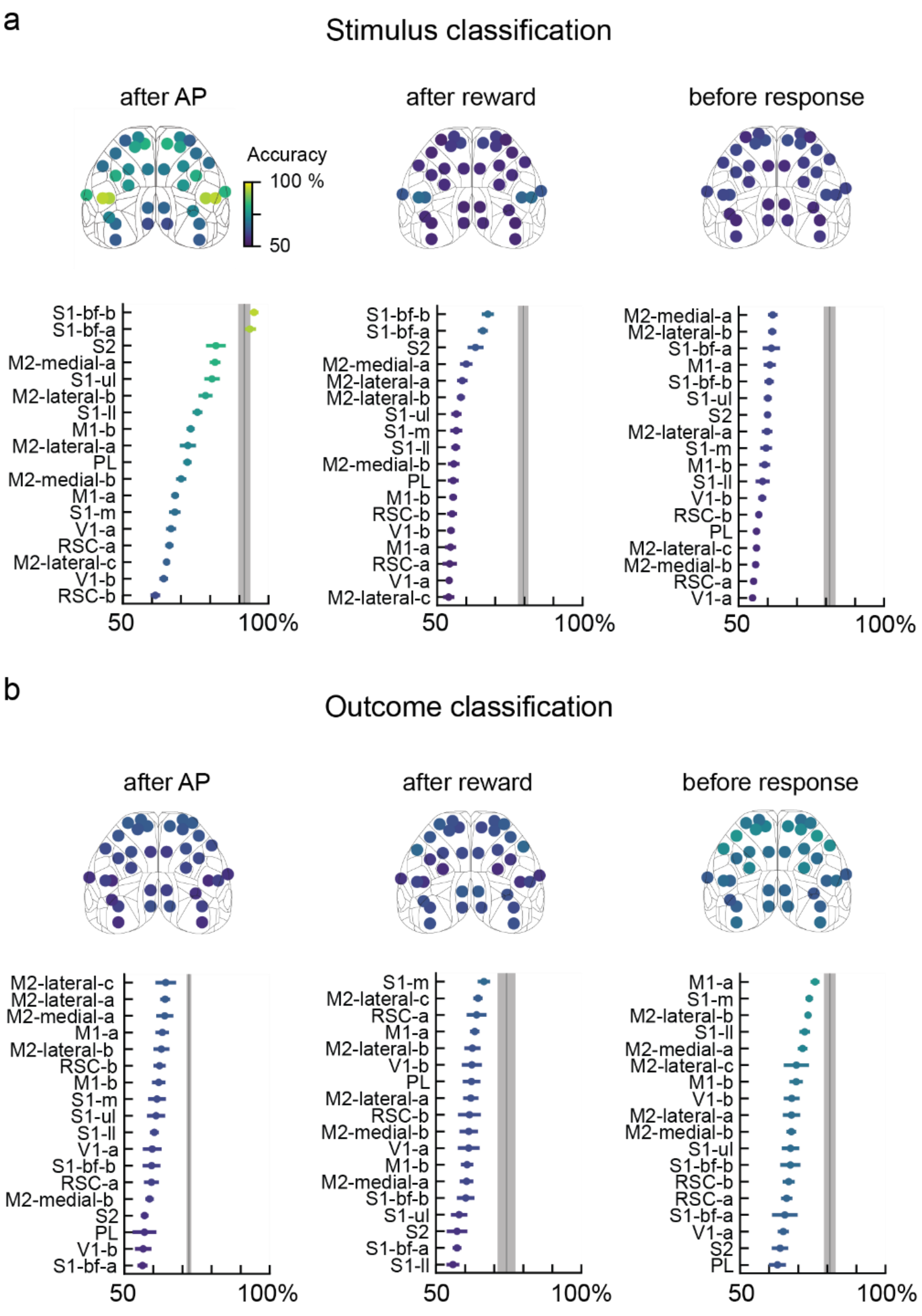
a
Stimulus classification
after AP
after reward
before response
Accuracy
100 %
50
S1-bf-b
S1-bf-a
S2
M2-medial-a
S1-ul
M2-lateral-b
S1-ll
M1-b
M2-lateral-a
PL
M2-medial-b
M1-a
S1-m
V1-a
RSC-a
M2-lateral-c
V1-b
RSC-b
S1-bf-b
S1-bf-a
S2
M2-medial-a
M2-lateral-a
M2-lateral-b
S1-ul
S1-m
S1-ll
M2-medial-b
PL
M1-b
RSC-b
V1-b
M1-a
RSC-a
V1-a
M2-lateral-c
M2-medial-a
M2-lateral-b
S1-bf-a
M1-a
S1-bf-b
S1-ul
S2
M2-lateral-a
S1-m
M1-b
S1-ll
V1-b
RSC-b
PL
M2-lateral-c
M2-medial-b
RSC-a
V1-a
50
100%
b
Outcome classification
after AP
after reward
before response
M2-lateral-c
M2-lateral-a
M2-medial-a
M1-a
M2-lateral-b
RSC-b
M1-b
S1-m
S1-ul
S1-ll
V1-a
S1-bf-b
RSC-a
M2-medial-b
S2
PL
V1-b
S1-bf-a
S1-m
M2-lateral-c
RSC-a
M1-a
M2-lateral-b
V1-b
PL
M2-lateral-a
RSC-b
M2-medial-b
V1-a
M1-b
M2-medial-a
S1-bf-b
S1-ul
S2
S1-bf-a
S1-ll
M1-a
S1-m
M2-lateral-b
S1-ll
M2-medial-a
M2-lateral-c
M1-b
V1-b
M2-lateral-a
M2-medial-b
S1-ul
S1-bf-b
RSC-b
RSC-a
S1-bf-a
V1-a
S2
PL
50
100%

**Supplementary Figure 3.** Behavioral classification from localized neural activity.

(a) Stimulus classification accuracies after AP (left), after reward (middle), and before response (right). Neural networks were given 200 ms windows of activity from left/right ROI pairs. Top colormaps show mean accuracies from corresponding ROI positions. Bottom plots show ROI accuracies from highest to lowest accuracy. Error bars indicate sem over 7 animals and gray shade shows mean ± sem of the prediction using all 18 ROI pairs (shown in Figure 6).

(b) Same as (a), but for outcome classification.

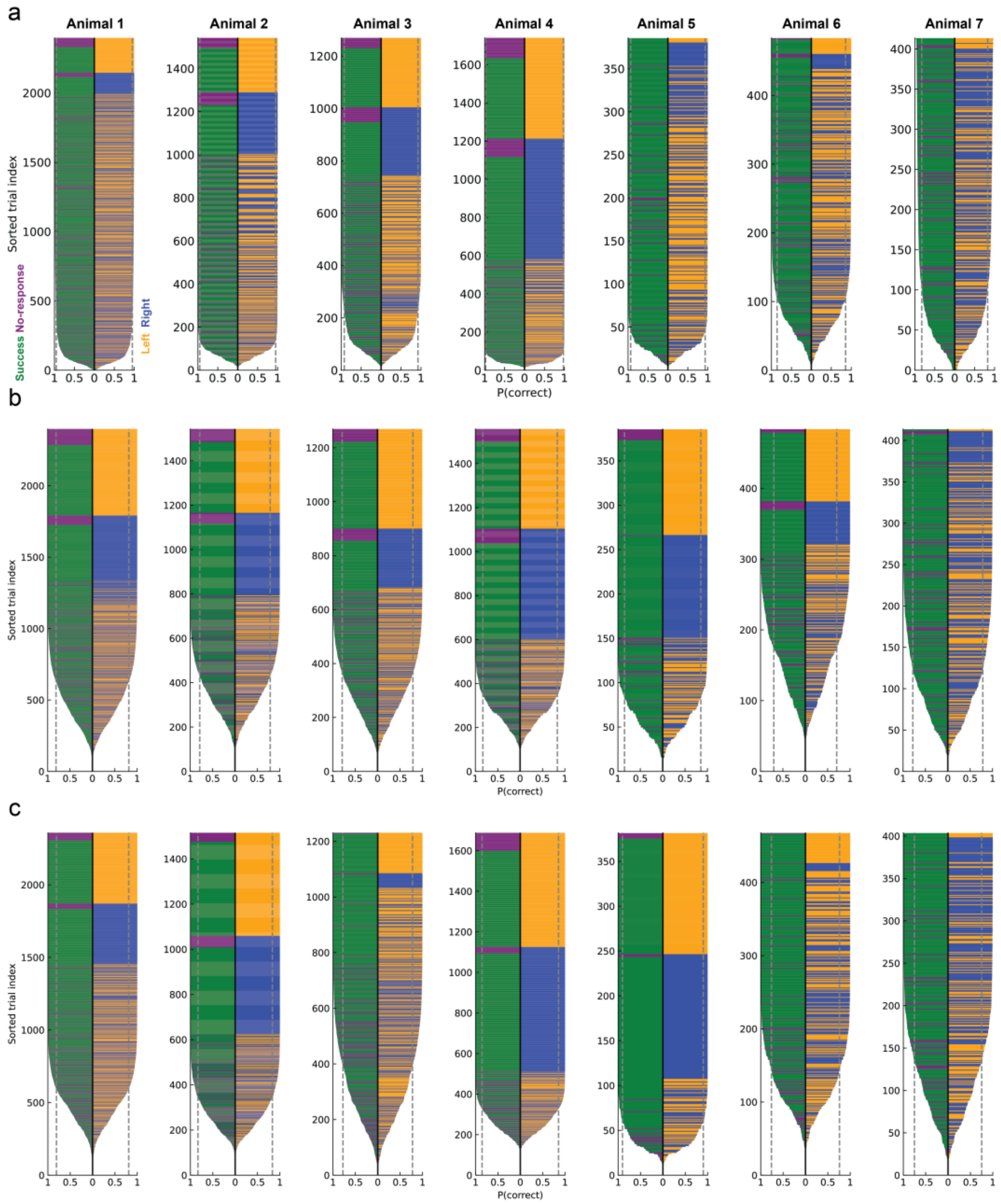


**Supplementary Figure 4.** Distribution of correct classification probabilities for all seven mice (columns) for stimulus classification using the $-\Delta F/F$ signal over a 200 ms period (a) after air-puff presentation, (b) after reward presentation, and (c) before response. Each horizontal line represents a trial whose length indicates the probability that it was classified correctly when in the test set (*N* = 800 model realizations on average for each trial; see Methods). The color of the left side of the line shows outcome (green if success, purple if no response) and that of the right shows stimulus (orange if left, blue right). The trials are sorted first according to the probability of correct classification, then according to stimulus, then outcome. Dotted line shows average accuracy.

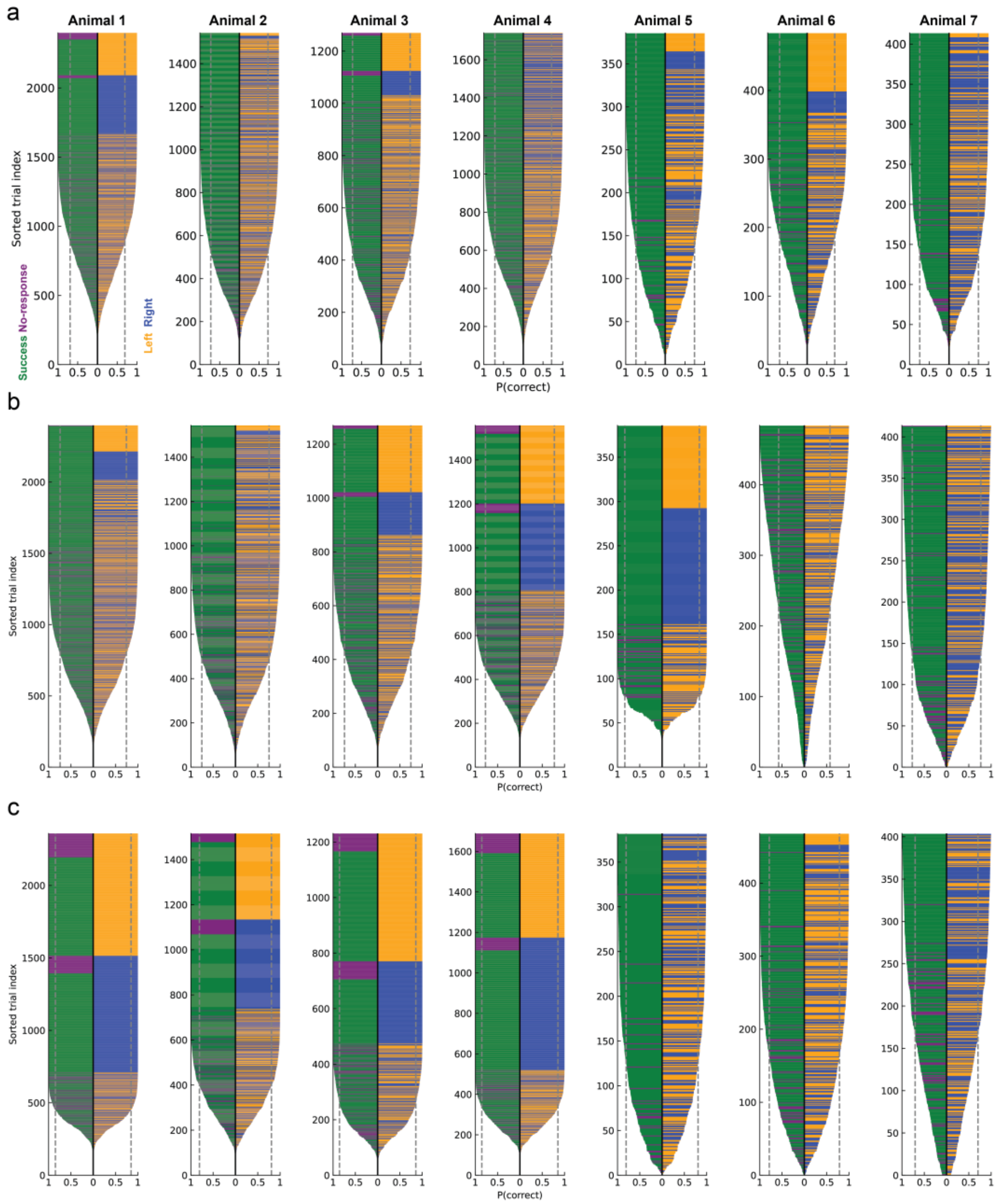


**Supplementary Figure 5.** Distribution of correct classification probabilities for all seven mice (columns) for outcome classification using the $-\Delta F/F$ signal over a 200 ms period (a) after air-puff presentation, (b) after reward presentation, and (c) before response. See Figure S4 for color legend. The mice that had worse outcome predictability later in the trial (row c, 3 rightmost panels) were those mice that had the reversed stimulus side vs. rewarded side task contingency (see Methods).

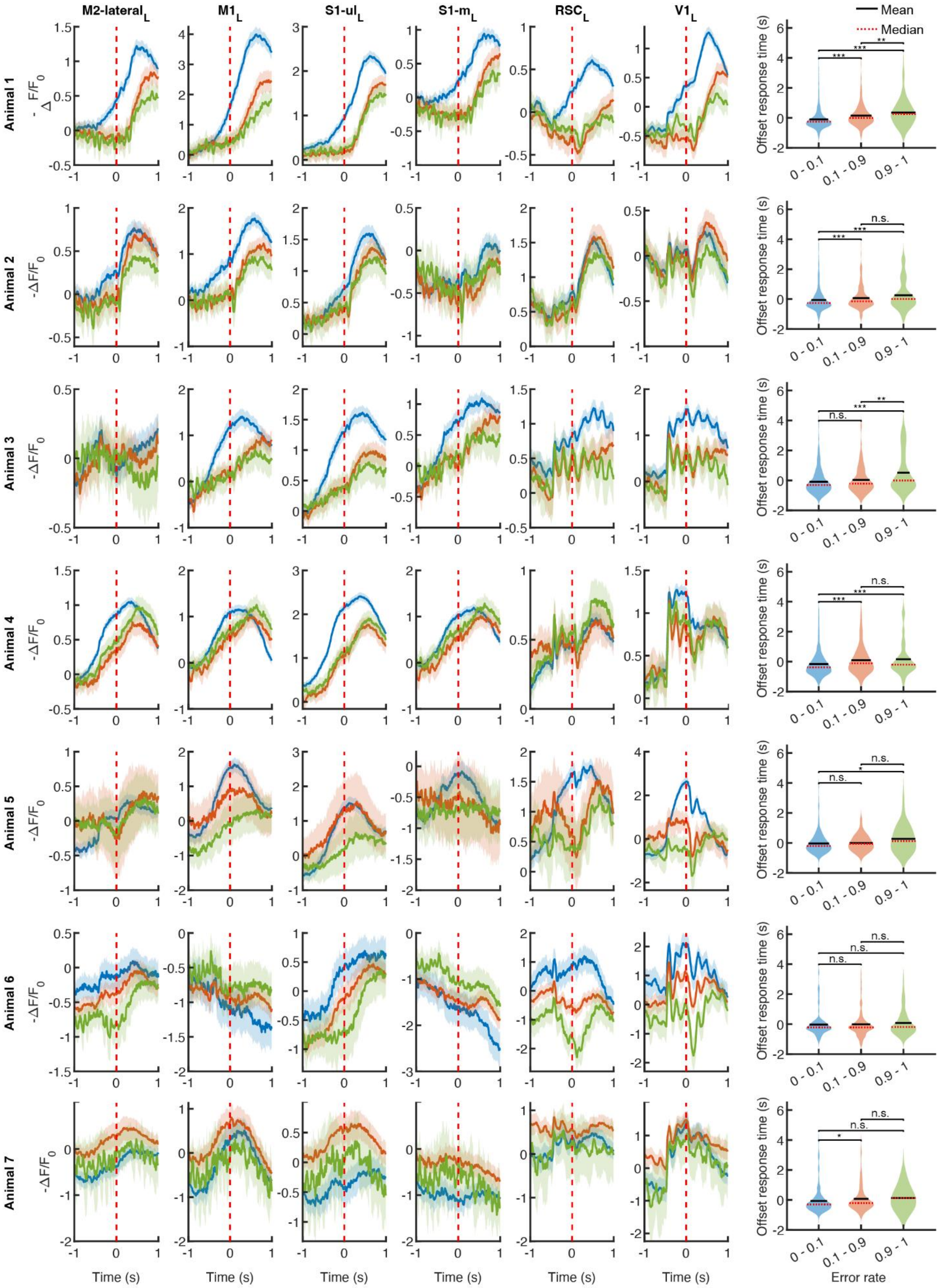

M2-lateral$_L$
M1$_L$
S1-ul$_L$
S1-m$_L$
RSC$_L$
V1$_L$
Mean
Median
Animal 1
Animal 2
Animal 3
Animal 4
Animal 5
Animal 6
Animal 7
-ΔF/F$_0$
Offset response time (s)
0 - 0.1
0.1 - 0.9
0.9 - 1
Time (s)
Error rate
n.s.

**Supplementary Figure 6.** Neural activity and response time are significantly different among groups with different error rates for outcome prediction in four out of seven animals. Each row shows $-\Delta F/F$ and response time of grouped trials based on the error rates. The first five columns show average neural activity at six ROIs from the left hemisphere. The right most column shows the distribution of response time among the three error rate groups. Lines denote mean $-\Delta F/F$ activity, and shaded areas denote 95% CI. P values for each animal are: (1) $4.5 \times 10^{-18}$, $4.2 \times 10^{-16}$, and 0.0032; (2) $4.3 \times 10^{-4}$, $6.6 \times 10^{-4}$, and 0.16; (3) 0.06, $2.6 \times 10^{-15}$, and 0.0048; (4) $5.9 \times 10^{-5}$, $9{,}3 \times 10^{-5}$, and 0.89; (5) 0.43, 0.05, and 0.33; (6) 0.98, 0.72, and 0.57; (7) 0.04, 0.27, and 0.63, all with Two-sided Mann-Whitney U Test. The animals with non-significant different response times and neural activity in motor cortex (bottom 3 rows) were in fact those animals with overall poorer outcome predictability (right 3 panels in Supp. Fig. 5) that were trained on the reverse stimulus side vs. rewarded side task contingency (see Methods).
